# Hypoxia threatens coral and sea anemone early life stages

**DOI:** 10.1101/2024.09.28.615579

**Authors:** Benjamin H. Glass, Katie L. Barott

**Affiliations:** Department of Biology, University of Pennsylvania, Philadelphia, PA 19104, USA

**Author notes:** **Contact** Benjamin H. Glass. Corresponding author: Katie L. Barott.

## Abstract

Seawater hypoxia is increasing globally and can drive declines in organismal performance across a wide range of marine taxa. However, the effects of hypoxia on early life stages (e.g., larvae and juveniles) are largely unknown, and it is unclear how evolutionary and life histories may influence these outcomes. Here, we addressed this question by comparing hypoxia responses across early life stages of three cnidarian species representing a range of life histories: the reef-building coral *Galaxea fascicularis*, a broadcast spawner with horizontal transmission of endosymbiotic algae (family Symbiodiniaceae); the reef-building coral *Porites astreoides*, a brooder with vertical endosymbiont transmission; and the estuarine sea anemone *Nematostella vectensis*, a non-symbiotic broadcast spawner. Transient exposure of larvae to hypoxia (dissolved oxygen < 2 mg L^-1^ for 6 h) led to decreased larval swimming and growth for all three species, which resulted in impaired settlement for the corals. Coral-specific responses also included larval swelling, depressed respiration rates, and decreases in symbiont densities and function. These results indicate both immediate and latent negative effects of hypoxia on cnidarian physiology and coral-algal mutualisms specifically. In addition, *G. fascicularis* and *P. astreoides* were sensitized to heat stress following hypoxia exposure, suggesting that the combinatorial nature of climate stressors will lead to declining performance for corals. However, sensitization to heat stress was not observed in *N. vectensis* exposed to hypoxia, suggesting that this species may be more resilient to combined stressors. Overall, these results emphasize the importance of reducing anthropogenic carbon emissions to limit further ocean deoxygenation and warming.

## Introduction

Dissolved oxygen (DO) has been deemed a “universal currency” in coastal marine ecosystems due to its fundamental role in aerobic respiration and dynamic cycling between organisms and the environment (Nelson and Altieri 2019). Seawater DO concentrations in coastal habitats can fluctuate between normoxia (≥ 5 mg DO L^-1^) and severe hypoxia (< 2 mg DO L^-1^; (Rosenberg 1980; Pezner et al. 2023)), with variability occurring over spatial and temporal scales (Altieri et al. 2017; Breitburg et al. 2018; Pezner et al. 2023). For example, daily troughs in DO on coral reefs occur overnight and can pass the threshold of hypoxia for several hours (Pezner et al. 2023), particularly during warm summer months when the solubility of molecular oxygen (O_2_) is lowest (Altieri et al. 2017; Hughes et al. 2020). Periods of low DO concentrations (i.e., hypoxic events) also occur seasonally on shallow reef habitats with limited flow (e.g., reef flats), where aerobic respiration of reef organisms consumes O_2_ that is not readily replenished by incoming oceanic seawater (Altieri et al. 2017; Hughes et al. 2020). In addition, hypoxia is common along continental margins, where terrestrial nutrient input and the upwelling of deep water can lead to transient (hours to days) or even permanent localized hypoxic zones (Summers 2001; Breitburg et al. 2018). Under environmental hypoxia, the depletion of oxygen in the diffusive boundary layer surrounding aquatic organisms can further increase the severity of tissue-level hypoxia (Shick 1990; Wangpraseurt et al. 2012; Linsmayer et al. 2020; Dellaert and Putnam 2023).

In response to historical variability in seawater DO conditions, coastal marine invertebrates have evolved a variety of mechanisms that promote hypoxia tolerance (Vaquer-Sunyer and Duarte 2011; Nelson and Altieri 2019; Altieri et al. 2021). Temperate cnidarians are among the most resilient taxa (Vaquer-Sunyer and Duarte 2011). For example, the benthic sea anemone *Nematostella vectensis* (class Hexacorallia) is able to persist in shallow coastal estuaries that may experience frequent and severe hypoxic events (Diaz et al. 1992; Summers 2001; Kiddon et al. 2003; Baumann et al. 2015), though the mechanisms of resilience in this species remain unexplored. Despite being in the same class as *N. vectensis*, tropical reef-building corals are more sensitive to hypoxia (Altieri et al. 2017, 2021; Nelson and Altieri 2019; Hughes et al. 2020). However, some coral species can maintain aerobic respiration and other key metabolic processes under hypoxic conditions (Johnson et al. 2021b; Dilernia et al. 2024), indicating some level of resilience against this stressor. While these responses may facilitate survival under hypoxia in cnidarian adults, it is unclear whether they are sufficient to protect early life stages, which are sensitive to abiotic stressors in these and other marine invertebrate taxa (Przeslawski et al. 2015; Foo and Byrne 2017). Thus, it is critical to compare early life hypoxia responses in temperate and tropical cnidarians displaying variation in evolutionary and life histories to better predict species’ trajectories.

Cnidarians display a variety of reproductive strategies, with some species that are gonochoric (i.e., separate sexes), hermaphroditic, or both. Further, species either carry out sexual reproduction via broadcast spawning (i.e., release of gametes into the water column) or brooding (i.e., internal fertilization and release of fully developed larvae; (Baird et al. 2009; Harrison 2011)). Cnidarian larvae are nearly always planktonic planulae; once fully developed, they must navigate the pelagic environment to locate suitable benthic substrates for settlement and metamorphosis into sessile juveniles (Baird et al. 2009; Harrison 2011). As in a broad diversity of animals, O_2_ is likely a central regulator of cnidarian development due to its roles in aerobic respiration, cell signaling, (epi)genetic regulation, and other processes (Alderdice et al. 2022b). Furthermore, the successful completion of cnidarian early life history stages hinges upon O_2_-dependent biological processes, such as ciliary motion and cellular respiration (Ricardo et al. 2016; Kitchen et al. 2020). These processes are sensitive to hypoxia in adult cnidarians (Murphy and Richmond 2016; Alva García et al. 2022; Pontes et al. 2023), and while transcriptomic and behavioral evidence suggests that larvae are similarly harmed (Jorissen and Nugues 2021; Alderdice et al. 2022b; Mallon et al. 2023), physiological data from a diversity of species are critically needed to better characterize these responses. In particular, comparing the effects of hypoxia on early life stages from cnidarians displaying variation in sexual system (i.e., gonochoric vs. hermaphroditic) and reproductive mode (i.e., spawner vs. brooder) is needed to better predict outcomes in future seas.

In addition to the direct influence on animal physiology during cnidarian development, hypoxia may also impact the performance of dinoflagellate algal endosymbionts (family Symbiodiniaceae). These symbionts are either passed to larvae through eggs (i.e., vertical transmission) or acquired *de novo* from the environment (i.e., horizontal transmission) during the larval or juvenile stages (Baird et al. 2009; Harrison 2011), and their photosynthetic activity provides fixed carbon to the cnidarian host, supplying a majority of the host’s nutrition (Jacobovitz et al. 2023; Helgoe et al. 2024). This mutualism contributes to variable O_2_ dynamics within host tissues through the consumption and production of O_2_ via respiration and photosynthesis, respectively (Rands et al. 1992; Linsmayer et al. 2020, 2024). These diel fluctuations in intracellular O_2_ production and consumption by symbionts can influence the holobiont (i.e., coral host and symbionts) response to seawater hypoxia (Kvitt et al. 2022; Helgoe et al. 2024). For example, symbiont photosynthesis results in the elevation of tissue O_2_ concentrations (Rands et al. 1992), which can help the host maintain performance in hypoxic seawater (Malcolm and Brown 1977). Further, hypoxia exposure can lead to a decline in symbiont photophysiology, the production of harmful reactive oxygen species (ROS) by symbiont photosynthetic machinery, and/or the loss of symbionts (i.e., bleaching) from coral tissue (Helgoe et al. 2024). The influence of hypoxia on symbiotic larvae and juveniles remains poorly understood, but declines in symbiont function under hypoxia would be expected to reduce host performance. Thus, it is vital to compare the hypoxia responses of cnidarian species that do and do not host symbionts.

Overall, the effects of hypoxia on cnidarian early life stages are largely unknown, and it is unclear how evolutionary and life histories may influence these outcomes. To address these key questions, we compared hypoxia responses across the early life stages of three cnidarian species: the sea anemone *N. vectensis*, a gonochoric and non-symbiotic spawner (Hand and Uhlinger 1992); the reef-building coral *Galaxea fascicularis*, a gonochoric spawner with horizontal symbiont transmission (Wei et al. 2023); and the coral *Porites astreoides*, a hermaphroditic brooder with vertical symbiont transmission (Goodbody-Gringley et al. 2018). We hypothesized that *N. vectensis* would generally be more tolerant of hypoxia than the corals due to this species’ adaptation to high DO variability and seasonal hypoxia in its native estuarine habitats (Vaquer-Sunyer and Duarte 2011). We also hypothesized that the non-symbiotic larvae of *N. vectensis* and *G. fascicularis* would be less severely impacted by hypoxia compared to *P. astreoides* due to use of cellular O_2_ by symbionts under hypoxia in the latter. However, we further hypothesized that *G. fascicularis* would be more sensitive to interactive hypoxia and heat stress compared to *P. astreoides*, as *Porites* spp. corals are generally more heat tolerant than other coral species (DeCarlo et al. 2019; Brown et al. 2023). To investigate these and other hypotheses, stage-matched larvae of each species were exposed to hypoxia for 6 h overnight, simulating an acute hypoxic event of the intensity and duration expected to become increasingly frequent in reefs and estuaries (Breitburg et al. 2018; Pezner et al. 2023; Rose et al. 2024). A suite of physiological, behavioral, and developmental traits was then assessed through the larval and into the juvenile stages. Overall, this study provides foundational knowledge of the effects of hypoxia on cnidarian early life stages.

## Materials and methods

### Adult culture and spawning

Full details regarding adult culture and spawning are in the supplementary material. Adult *Nematostella vectensis* sea anemones were collected from a salt marsh (Brigantine, NJ), and spawning was induced by exposing anemones (*n* = 200) to light and elevated temperatures (24°C) for 14 h, followed by transfer to ∼18–19°C (Hand and Uhlinger 1992). Larvae (*n* = 1 cohort with mixed parentage) were then cultured to the planula stage (3 days post-fertilization) for experimentation. An aquarium population of adult *Galaxea fascicularis* colonies (*n* = 9 females, 10 males) spawned during August 2023 in an *ex situ* system (Craggs et al. 2017) located at Carnegie Science (Baltimore, MD, USA), yielding a cohort of planulae (*n* = 1 cohort with mixed parentage) that were used in experiments within 48 h. Adult *Porites astreoides* colones (*n* = 20) were collected in Bermuda (32°22’13” N, 64°44’27” W) during July 2023 and maintained in aquaria under ambient conditions (Wong et al. 2021). Brooded planulae (*n* = 4 cohorts) were collected according to previously developed procedures (Goodbody-Gringley et al. 2018), and maintained in culture for ∼48 h. At the time of use in experimentation, larvae from all 3 species were in the planula stage and would become competent to settle within 72–96 h (Hand and Uhlinger 1992; Goodbody-Gringley et al. 2018; Wei et al. 2023). Artificial seawater was used for culturing and experimentation for *N. vectensis* and *G. fascicularis*, while flow-through, natural seawater facilities at the Bermuda Institute for Ocean Science were used for *P. astreoides*.

### DO treatments

Methods for the DO treatments were consistent across the three species. We chose 1.6 mg DO L^-1^ as our hypoxia DO concentration and 6 h as the length of the treatment period in accordance with current and expected future conditions on coral reefs and in estuaries (Summers 2001; Breitburg et al. 2018; Pezner et al. 2023). As troughs in seawater DO typically occur at night (Summers 2001; Pezner et al. 2023), treatments were performed from 21:00–3:00, with red light headlamps as the only source of illumination. Shortly before 21:00, stage-matched, swimming planulae (*n* = 1200–2400 larvae cohort^-1^ species^-1^) were distributed with 1–2 mL of seawater across six, 15 mL conicals representing three replicate subgroups for each of the two treatments (normoxia/control and hypoxia; *n* = 600–1200 larvae treatment^-1^ cohort^-1^ species^-1^). Prior to the treatment, 20–30 larvae from each subgroup (*n* = 60–90 larvae treatment^-1^ cohort^-1^ species^-1^) were transferred to a 6-well plate and photographed under a dissecting microscope for size measurements. Next, a graduated cylinder (2 L) was filled with seawater and the temperature (YSI ProSolo; accuracy = ± 0.2°C, precision = 0.1°C) and salinity (accuracy = ± 0.1 ppt, precision = 0.01 ppt) were recorded. This seawater was used to fill 3 glass jars (500 mL). A DO probe (YSI ProSolo; accuracy = ± 0.1 mg L^-1^, precision = 0.01 mg L^-1^) was submerged in each jar and nitrogen gas was bubbled through an airstone into the seawater until the target DO (1.6 mg L^-1^) was reached. Larvae were then poured into the jars, and final DO adjustments were made before the jars were sealed and placed at each species’ ambient culture temperatures (*N. vectensis*: 18°C; *G. fascicularis*: 27°C; *P. astreoides*: 28°C) using either incubators (for *N. vectensis* and *G. fascicularis*; Bokel Scientific) or a seawater bath (for *P. astreoides*) for 6 h. The remaining larvae were distributed into three additional jars with seawater that was not deoxygenated (i.e., normoxia controls). DO measurements were again collected, and these jars were also sealed and placed alongside the hypoxia jars for 6 h. While deoxygenation via N_2_ gas can result in relatively small increases in pH due to the simultaneous elimination of CO_2_ (Johnson et al. 2021b), the pH of the jars was not measured, which is an important consideration for interpreting the results.

After 6 h, each jar was uncovered and an image was collected using an iPhone 14 (Apple) with the plane of focus at the bottom of the jar. Within seconds, a DO measurement was also collected. Later, the jar images were used to determine the percentages of larvae in each jar actively swimming in the water column as follows: (1-(larvae at bottom/total larvae))*100. Next, all larvae were transferred to culture containers with new (normoxic) seawater. For *N. vectensis*, larvae were cultured inside a dark incubator (18°C) in Petri dishes with ∼25 mL of seawater, whereas coral larvae were cultured in 500 mL plastic tubs. For *G. fascicularis*, the tubs were placed in an incubator (27°C) under a light set to a 12-h:12-h day:night cycle (∼50 µmol quanta m^-2^ s^-1^), and the water was changed daily. Containers housing *P. astreoides* larvae were modified to add mesh holes in the sides, and were placed on tables of egg crate in a flow-through system under ambient reef temperature (∼28°C) and light conditions as described above. Culture conditions represent ambient conditions for each species, and match the conditions experienced by the adults from which gametes were sourced prior to spawning.

### Larval in vivo metrics and sampling

Sampling procedures for larvae were performed immediately following the treatment period (0 h post-treatment), as well as at 12 and 36 h post-treatment. To assay the photosystem II photochemical yield (F_v_/F_m_) of *P. astreoides* larval endosymbionts, 10 larvae from each subgroup (*n* = 30 larvae treatment^-1^ time point^-1^ cohort^-1^) were first transferred to a 6-well plate, which was placed in a dark incubator (Marshall Scientific) at ∼28°C. The larvae were dark-acclimated for 30 min, then transferred in 80 µL of seawater to the tip of a diving pulse-amplitude-modulation (PAM) fluorometer (Heinz Walz). A single measurement (gain = 4, damping = 18, ERT-factor = 0.73, measuring light intensity = 7, measuring light frequency = 3, SAT intensity = 8, SAT widths = 0.6) was then collected. To determine larval size (length in mm), 20–30 larvae from each subgroup (*n* = 60–90 larvae treatment^-1^ time point^-1^ cohort^-1^ species^-1^) were transferred to a 6-well plate and photographed under a dissecting microscope (AmScope). A ruler was also photographed under identical microscope settings. Images were later analyzed in FIJI (Schindelin et al. 2012). Specifically, the ruler image was used to set the scale, and then the line tool was used to measure the maximum length (i.e., major axis) of each larva. The width of each larva was also measured and used to calculate surface area, volume, and SA:V ratios using equations for an elliptical (prolate) sphere (de Putron et al. 2017).

Larval heat tolerance was assayed using previously established methods for coral and sea anemone larvae with modifications (Glass et al. 2023b; Zhang et al. 2023b). Specifically, 10 larvae from each subgroup (*n* = 30 larvae treatment^-1^ time point^-1^ cohort^-1^ species^-1^) were first transferred to a 1.5 mL tube with 1 mL of seawater (for *P. astreoides*) or to individual wells of a 96-well PCR plate with 100 µL of seawater (for *N. vectensis* and *G. fascicularis*). The larvae were then placed at 36°C (no ramp period) either in a water bath with a 50 W aquarium heater (Aqueon) or a PCR thermocycler (Thermo Fisher Scientific) for tubes and plates, respectively. This temperature was chosen based on methods developed for larvae of the coral *Orbicella faveolata* (Zhang et al. 2023b), and was intended to induce mortality over time. At several time points every 1–12 h, the number of surviving larvae (i.e., the number present in the tubes, as dead larvae quickly dissolve) was recorded. Data were collected until all larvae had experienced mortality (*N. vectensis*: ∼125 h; *G. fascicularis*: ∼50 h; *P. astreoides*: ∼75 h). For symbiotic *P. astreoides* larvae, photochemical yield was also assayed throughout the heat treatment using a diving PAM fluorometer as described.

To determine larval metabolic rates, three subgroups of 20 (for *G. fascicularis* and *P. astreoides*) or 30 (for *N. vectensis*) larvae were transferred to a 24-well plate (*n* = 1 well subgroup^-1^, 60–90 larvae treatment^-1^ time point^-1^ cohort^-1^ species^-1^) equipped with oxygen sensor spots (Loligo Systems). All wells were filled to capacity (80 μL) with seawater, and 3–4 wells were also filled with culture seawater to serve as controls. The plate was sealed with an adhesive plate cover and placed on a calibrated PreSens SensorDish Reader (Precision Sensing) in a dark incubator at the appropriate culture temperature for each species (*N. vectensis*: 18°C; *G. fascicularis*: 27°C; *P. astreoides*: 28°C). The oxygen concentration (μmol O_2_ L^-1^; accuracy = ± 1 µmol, precision = 0.01 µmol) in each well was recorded every 15 s for 1 h, during which oxygen consumption rates remained relatively constant, indicating a lack of oxygen limitation. To determine photosynthesis rates of *P. astreoides* endosymbionts, a 16 HD LED reef light (AquaIllumination) set to achieve a photosynthetically active radiation (PAR) level of ∼200 µmol m^-2^ s^-1^ (Apogee Instruments; accuracy = ± 5%, precision= 1 µmol m^-2^ s^-1^) was placed above the plate following respiration measurements, and the oxygen concentration (μmol O_2_ L^-1^) in each well was again recorded every 15 s for 1 h. Following data collection, the larvae in each well (*n* = 60–90 larvae treatment^-1^ time point^-1^ cohort^-1^ species^-1^) were transferred to 1.5 mL tubes. Seawater was removed from the tubes, which were then stored at -80°C for later determination of additional physiological metrics (see below). The slopes of best-fit linear regressions relating the DO concentration (µmol O_2_ L^-1^) in each well to time (in min) were generated. These rates were then converted to pmol O_2_ consumed (respiration) or produced (photosynthesis) per min, expressed as positive values. Later, rates for each well were normalized to the organic biomass (ash-free dry weight in µg; see below) in the same well.

### Juvenile in vivo metrics and sampling

Following the 36 h larval sampling, remaining *N. vectensis* and *G. fascicularis* larvae (*n* = 300–390 larvae treatment^-1^ time point^-1^ cohort^-1^ species^-1^) were maintained in their culture dishes for settlement. The number of settled juveniles was counted to determine the settlement rate (%) in each dish daily until all larvae either settled or experienced mortality. To quantify juvenile size (polyp diameter in mm), images of juveniles were collected at 84 h post-treatment and analyzed in FIJI as previously described. For later quantification of physiological metrics, 20 (for *G. fascicularis* and *P. astreoides*) or 30 (for *N. vectensis*) juveniles from each subgroup (*n* = 60–90 juveniles treatment^-1^ species^-1^) were also sampled at 84 h. For the corals, a razor blade or cut pipette tip was used to remove the juveniles from the walls of the plastic culture containers. Unattached juveniles for both species were transferred to 1.5 mL tubes, which were stored at -80°C until further analysis. Symbiosis establishment was assayed in aposymbiotic *G. fascicularis* juveniles (see supplementary materials for full details). Briefly, Symbiodiniaceae (∼500,000 cells per culture container; taxonomic composition unknown) were supplied to juveniles shortly after 84 h, which fell during the competence window for uptake (Wei et al. 2023). After 6 h, water in culture containers was stirred; after an additional 18 h, a water change was performed and the symbiotic juveniles were sampled (*n* = 15–30 juveniles treatment^-1^) by culture container and stored at -80°C for later analysis of symbiont density (see below).

For *P. astreoides* cohorts 1–3, larvae remaining after the 36 h sampling (*n* = 210–450 larvae treatment^-1^ cohort^-1^) were induced to settle as previously described (Goodbody-Gringley et al. 2018). Specifically, larvae were transferred to 500 mL plastic containers (*n* = 1 container treatment subgroup^-1^) with mesh tops (150 µm), which allowed for water flow while keeping larvae inside. Each container was equipped with egg crate supporting five terracotta tiles (25 cm^2^) that had been preconditioned at a depth of ∼3 m on the reef for ∼6 mo. The containers were placed at ambient temperatures at the bottom of flow-through tanks. After 48 h (i.e., at 84 h post-treatment), the containers were uncovered and the settled juveniles in each were counted and imaged under a dissecting microscope along with a ruler. These images were later used to qualify juvenile size (polyp diameter in mm) using FIJI as previously described. Following imaging, 20 juveniles from each subgroup (*n* = 60 juveniles treatment^-1^ cohort^-1^) were sampled for physiology (see below). Specifically, a razor blade or cut pipette tip was used to remove the juveniles from the settlement tiles, followed by transfer to 1.5 mL tubes. Seawater was removed from the tubes, which were then stored at -80°C. For one additional cohort of *P. astreoides* larvae (*n* = 380 larvae treatment^-1^), animals were maintained in culture containers until 84 h, after which settlement in the water column was observed, allowing for nondestructive manipulation of juveniles. Settlement rates for these containers were quantified, and then the juveniles were assayed for photochemical yield and metabolic rates before storage at -80°C.

### Physiology sample processing

Larval and juvenile physiology samples for all three species were processed in parallel. The samples were first put on ice, and 200 μL of chilled 1x tris-NaCl-EDTA lysis buffer supplemented with 1 mM dithiothreitol, protease inhibitor cocktail (Thermo Fisher Scientific), and phosphatase inhibitors (Roche), were added to each. Cell lysis was achieved via vigorous pipetting followed by sonication (Diagenode UCD-200) at 4°C for 5 min with a 30:60 s on:off cycle. For samples containing symbionts (i.e., all *P. astreoides* samples and *G. fascicularis* juveniles following symbiont infection), tissue homogenates were centrifuged at 2,500 x *g* for 5 min at 4°C, then the supernatants (host slurry) were transferred to new tubes while the pellets (symbionts) were stored on ice. For all species, host slurry was then aliquoted in equal volumes into two, 1.5 mL tubes. Next, one of the host slurry aliquots was used for protein determination (Whitaker and Granum 1980). Specifically, 25 µL of each sample were pipetted in triplicate into wells of a UV-transparent 96-well plate (Thermo Fisher Scientific). Three wells containing 25 µL of lysis buffer were also included as blanks. Next, 175 µL of 1x phosphate buffered saline (PBS) were added to each well, and the absorbance of the wells at 235 and 280 nm was measured using a spectrophotometer (Agilent). All absorbance values were confirmed to be between 0.1 and 1, and then the average values for the blanks were subtracted from the values for each well. Protein concentrations (mg L^-1^) were calculated by subtracting the blank-corrected absorbance of each well at 280 nm from that of the same well at 235 nm and dividing by 2.51 (Whitaker and Granum 1980). Concentrations were then averaged across the three technical replicates, multiplied by the original animal slurry volume (200 µL) and divided by the average size (mm) of the animals in each sample.

Assays were performed to determine the ash-free dry weight (AFDW; i.e., organic biomass) of the animal tissue in each sample. Specifically, aluminum pans were labeled with pencil and combusted in a muffle furnace (Thermo Scientific Lindberg Blue M) at 450°C for 4 h. After cooling, 100 µL of host slurry was added to the pans, which were then placed in a drying oven (Quicy Lab, Inc. model 10 lab oven) at ∼60°C. Pans were dried until a constant weight was achieved (∼24 h), weighed on a microbalance (Sartorius CPA225D; accuracy = ± 0.00002 g, precision = 0.00001 g), and combusted at 450°C for 4 h before being weighed again. The AFDW of each sample was calculated by subtracting the weight of the dry tissue after combustion from that of the dry tissue before combustion, then normalized to size by dividing the AFDW (µg) by the average size (mm) of the animals in each sample. To determine symbiont densities, symbiont pellets were first resuspended in 150 µL of 0.1% SDS in filtered seawater. Next, 15 µL (for *P. astreoides* samples) or 50 µL (for *G. fascicularis* samples) of each sample were pipetted in triplicate into wells of a 96-well plate. Flow cytometry (Guava® easyCyte™ HT) was employed to determine the concentration of symbionts in each well using gates based on forward scatter, side scatter, and red (chlorophyll) autofluorescence excited at 488 nm (Krediet et al. 2015; Innis et al. 2021). The concentrations of symbionts in each well were corrected for volume, averaged across the replicates, and then normalized to AFDW.

To determine the chlorophyll content of *P. astreoides* symbionts (no symbionts remained after density determination for *G. fascicularis*), the remaining 105 µL of resuspended symbionts was first centrifuged at 2,500 x *g* for 5 min in a chilled (4°C) centrifuge to pellet symbionts, then the supernatant was removed and replaced with 125 µL of 100% acetone. Samples were thoroughly vortexed, then placed at -80°C for 24 h, after which vortexing was repeated and the samples were returned to -80°C for an additional 24 h. Following incubation, samples were vortexed again, then centrifuged at 2,500 x *g* for 5 min in a chilled (4°C) centrifuge to pellet the cells. Next, 100 µL of the supernatants (acetone containing extracted chlorophyll) from each tube were pipetted into wells of a 96-well plate, which was immediately brought to a spectrophotometer (Agilent) for absorbance readings at 630, 663, and 750 nm. The weight of chlorophyll (g) in each well was determined according to the following equation, which included a correction for the sample pathlength (0.29 cm): ((11.43*(A_663_-A_750_))-(0.64*(A_630_-A_750_)))/0.29 (Banse and Anderson 1967). These weights were then used in combination with symbiont densities to calculate chlorophyll density expressed as pg symbiont cell^-1^.

### Data and statistical analyses

Full details regarding data and statistical analyses are present in the supplementary material. RStudio with R version 4.2.1 was used for all analyses (RStudio Team 2020). First, survival and photochemical yield data from larval heat tolerance assays were used to create dose-response curves using the package *drc* (Ritz et al. 2015), and an LD50 was determined for each curve (*n* = 3 curves treatment^-1^ time point^-1^ cohort^-1^ species^-1^) using the package *chemCal* (Ranke 2022). Linear models were constructed relating each metric to the interaction between species, treatment, and/or time (h post-treatment) as appropriate, confirmed to meet relevant assumptions, and subjected to an analysis of variance (ANOVA) to determine the statistical significance of the model terms. Finally, Tukey’s Honest Significant Difference post-hoc tests were used to determine the significance of pairwise comparisons where appropriate. All values are expressed as averages rounded to appropriate significant figures ± standard error of the mean (SEM), and all original data and code are publicly available online (Glass and Barott 2024).

## Results

### Experimental conditions

Seawater temperatures and dissolved oxygen (DO) concentrations within the hypoxia (∼1.6 mg DO L^-1^) and normoxia/control (> 6.5 mg DO L^-1^) treatments (*n* = 3 groups per treatment) were relatively constant over the treatment period (6 h) for all three species (*Nematostella vectensis*, *Galaxea fascicularis*, and *Porites astreoides*; Figure 1a–b). For *N. vectensis*, the mean (± standard error [SEM]) temperature and DO concentration over the treatment period were 19.2 ± 0.1°C and 8.68 ± 0.01 mg L^-1^ (94 ± 0.1% saturation; normoxia), versus 19.1 ± 0.1°C and 1.59 ± 0.01 mg L^-1^ (17.2 ± 0.1%; hypoxia). For *G. fascicularis*, the temperature and DO concentration over the treatment period were 26.6 ± 0.2°C and 7.59 ± 0.01 mg L^-1^ (94.6 ± 0.1%; normoxia), versus 26.5 ± 0.2°C and 1.65 ± 0.01 mg L^-1^ (20.6 ± 0.1%; hypoxia). Finally, for *P. astreoides*, the temperature and DO concentration over the treatment period were 27.8 ± 0.2°C and 6.8 ± 0.1 mg L^-1^ (86.6 ± 1.3%; normoxia), versus 27.7 ± 0.2°C and 1.7 ± 0.1 mg L^-1^ (21.7 ± 1.3%; hypoxia). Mean (± SEM) temperatures and DO concentrations at the beginning and end of the treatment periods are reported for each species in Table S1.

**Figure 1.**
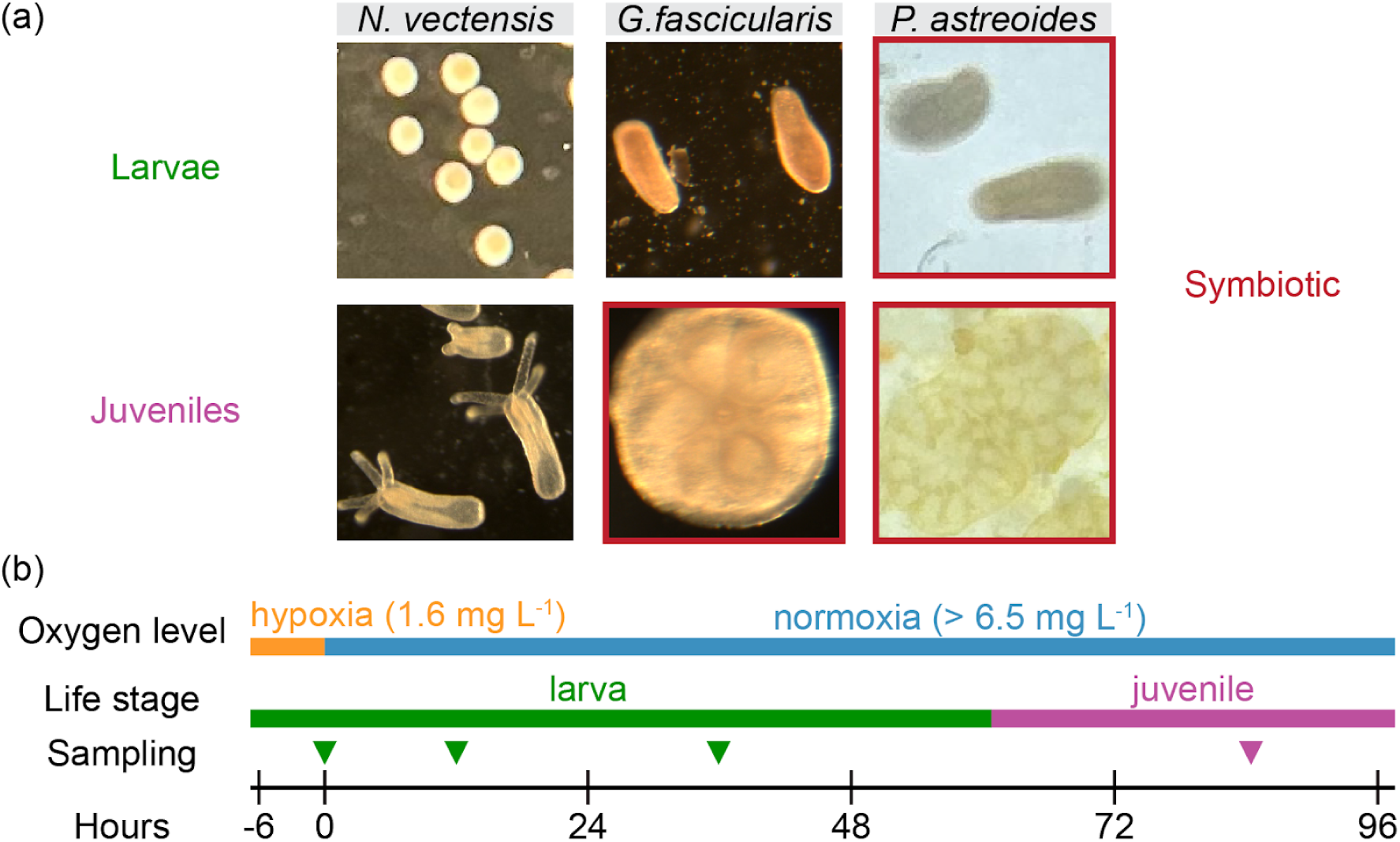
Study species and experimental timeline. (a) Representative images of larvae (top row) and juveniles (bottom row) from the three study species: *Nematostella vectensis* (left), *Galaxea fascicularis* (middle), and *Porites astreoides* (right). Red borders around images indicate life stages that possess dinoflagellate endosymbionts (family Symbiodiniaceae). (b) Timeline of the experiment depicting the dissolved oxygen level of seawater in which organisms were held during and after the hypoxia treatment (control animals experienced normoxia throughout the experiment), organism life stages (larva = green, juvenile = purple), and sampling time points (arrowheads) colored by life stage at the time of sampling. Hours are relative to the end of the oxygen treatments (i.e., h post-treatment).

### Effects of hypoxia on larval swimming behavior and settlement

For all metrics, pertinent statistical testing information is reported in Table S2, while detailed statistical outputs are available in the public, permanent repository created for this publication (Glass and Barott 2024). Exposure to hypoxia for 6 h had significant, negative effects on larval swimming behavior and settlement success compared to normoxia for all three species (Figure 2a–c). The percentage of larvae actively swimming in the water column at the conclusion of the treatment period was significantly influenced by species, DO treatment, and their interaction (Table S2). Significantly fewer larvae were observed actively swimming in the water column for *N. vectensis*, *G. fascicularis*, and *P. astreoides* following exposure to hypoxia (32 ± 4%, 57 ± 7%, and 51 ± 2%, respectively) compared to controls (54 ± 1%, 94 ± 2%, and 71 ± 1%, respectively), while the total percentage of larvae swimming differed between species (Figure 2a–c). Settlement rates (expressed as the percentage of larvae settled) were also affected by species, treatment, and their interaction (Table S2). For *N. vectensis*, no significant difference in final settlement rate was observed between larvae exposed to hypoxia (93 ± 1%) compared to controls (94 ± 2%; Figure 2a). By contrast, both *G. fascicularis* and *P. astreoides* displayed significantly lower settlement rates for larvae exposed to hypoxia (42 ± 5% and 13 ± 1%, respectively) compared to controls (69 ± 4% and 35 ± 2%, respectively; Figure 2b–c).

**Figure 2.**
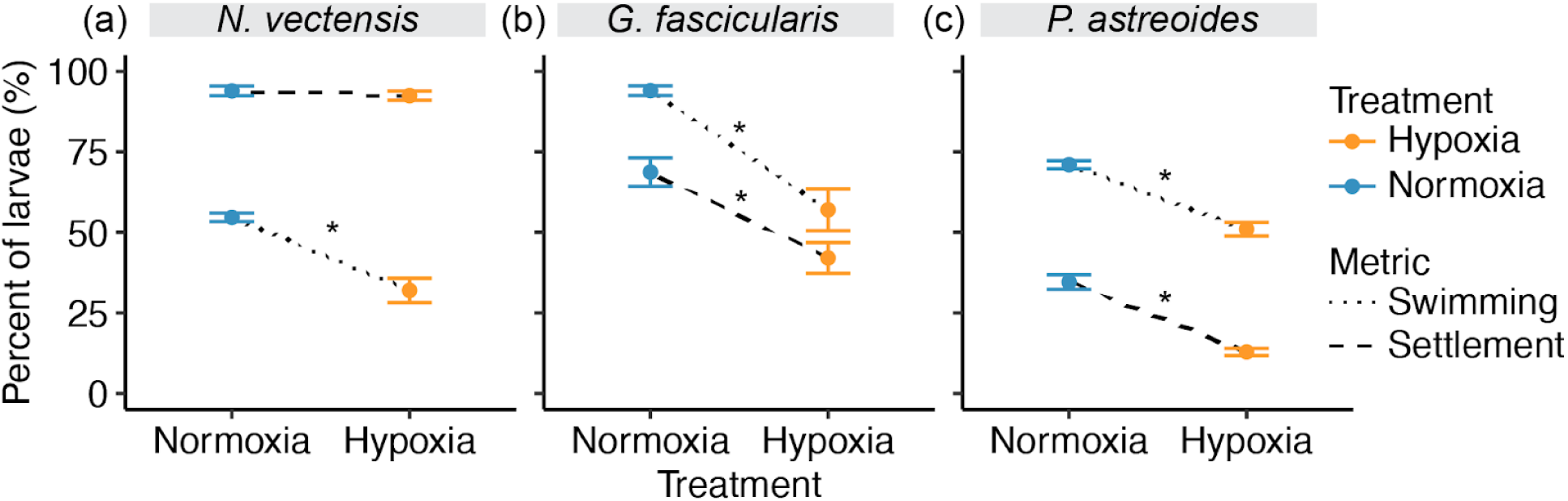
Effects of hypoxia on larval swimming and settlement. (a–c) Percent of (a) *Nematostella vectensis*, (b) *Galaxea fascicularis*, and (c) *Porites astreoides* larvae (*n* = 600–1200 larvae treatment^-1^ cohort^-1^ species^-1^) swimming at the conclusion of the 6-h treatment period (dotted lines) or settled (final percentage; dashed lines) for the hypoxia (yellow) and normoxia (blue) treatments. Points with error bars depict means ± SEM, and asterisks indicate statistical significance (*p* < 0.05) of pairwise comparisons (hypoxia vs. normoxia).

### Effects of hypoxia on animal growth and metabolism

Exposure to hypoxia had significant effects on larval and juvenile growth (i.e., size and biomass) and respiration rates compared to controls, with variable effect sizes and directions across species, life stages, and hours post-treatment (h; Figure 3a–i). Prior to the treatment period, larvae of all three species displayed no significant difference in size (length) between the treatment subgroups (*n* = 3 subgroups treatment^-1^; Figure S1a–c). Following the treatments, animal sizes (larval length and juvenile polyp diameter) were significantly influenced by species, treatment, h, and their interaction (Table S2). For *N. vectensis*, larvae exposed to hypoxia displayed no significant differences in size from controls at any time point (0.3–0.34 mm; Figure 3a). By contrast, larvae of both *G. fascicularis* (0.55–0.57 mm) and *P. astreoides* (0.71–0.73 mm) exposed to hypoxia were significantly larger than controls (*G. fascicularis*: 0.39–0.43 mm; *P. astreoides*: 0.56–0.64 mm) at all time points (Figure 3b–c). Interestingly, these trends reversed during the juvenile stage (84 h), at which point juveniles previously exposed to hypoxia were significantly smaller than controls for all species (*N. vectensis*: 0.21 ± 0.01 vs. 0.31 ± 0.01; *G. fascicularis*: 0.54 ± 0.01 vs. 0.67 ± 0.01; *P. astreoides*: 0.78 ± 0.02 vs. 0.9 ± 0.02 mm), including *N. vectensis*, which had not previously shown differences (Figure 3a–c). Larval width, surface area, and volume showed similar trends, but these data are not presented for simplicity and because length was considered analogous to juvenile polyp diameters.

**Figure 3.**
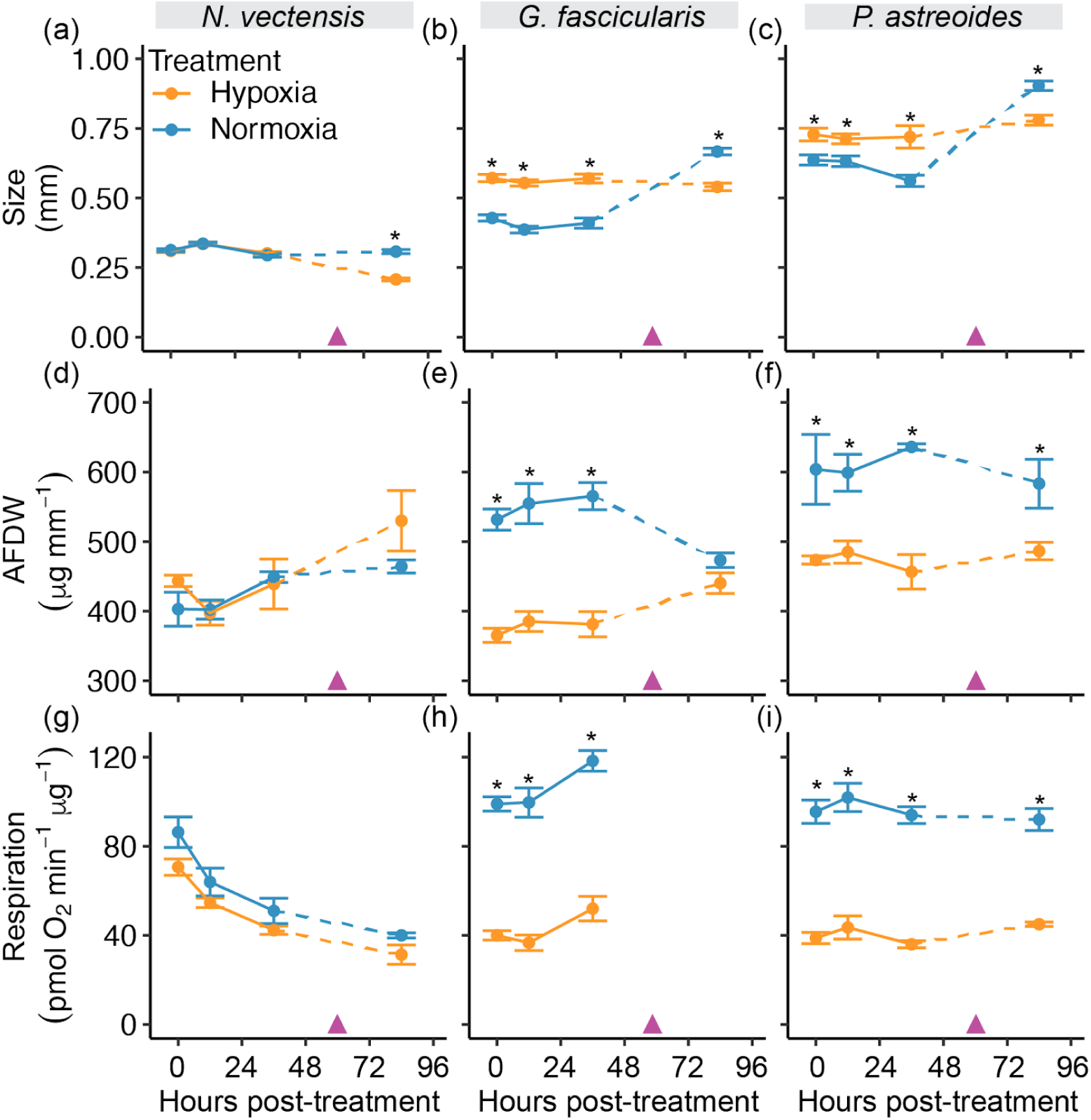
Effects of hypoxia on animal growth and metabolism. (a–c) Size (larval length or juvenile polyp diameter in mm), (d–f) ash-free dry weight (AFDW; normalized to size; µg mm^-1^), and (g–i) respiration rates (pmol O_2_ consumed min^-1^ µg AFDW^-1^) of *Nematostella vectensis* (left), *Galaxea fascicularis* (middle), and *Porites astreoides* (right) larvae and juveniles (*n* = 60–90 treatment^-1^ time point^-1^ cohort^-1^ species^-1^; dashed line separates life stages) over time following hypoxia (yellow) and normoxia (blue) treatments. Purple arrowheads indicate the approximate timing of settlement, points with error bars depict means ± SEM, and asterisks indicate statistical significance (*p* < 0.05) of pairwise comparisons (hypoxia vs. normoxia).

As with size, animal ash-free dry weight (AFDW; i.e., organic biomass) was significantly affected by hypoxia in a species-dependent matter. Specifically, AFDW (expressed normalized to size in µg mm^-1^) was significantly influenced by species, treatment, and their interaction. For *N. vectensis*, AFDW was statistically indistinguishable across treatments and life stages for animals exposed to hypoxia (397–530 µg mm^-1^) compared to controls (402–464 µg mm^-1^) at all time points (Figure 3d). By contrast, *G. fascicularis* larvae displayed significantly lower AFDW when exposed to hypoxia (365–385 µg mm^-1^) than controls (532–565 µg mm^-1^) at all time points (Figure 3e). At the juvenile stage, however, AFDW of *G. fascicularis* previously exposed to hypoxia were not significantly different from controls (440 ± 15 µg mm^-1^ vs. 473 ± 10 µg mm^-1^, respectively; Figure 3e). Interestingly, *P. astreoides* displayed a different trend, with animals exposed to hypoxia having significantly lower AFDW (457–486 µg mm^-1^) than controls (583–636 µg mm^-1^) at all time points (Figure 3f). Animal total protein (expressed normalized to size in µg mm^-1^) followed similar trends to AFDW (Figure S1d–f). As with the growth metrics quantified here (i.e., size, AFDW, and protein), respiration rates (expressed as pmol O_2_ consumed minute^-1^ µg AFDW^-1^) were significantly influenced by the interaction between species and treatment (Table S2). For *N. vectensis*, respiration rates were not significantly different for animals exposed to hypoxia (31–71 pmol O_2_ min^-1^ µg^-1^) compared to controls (40–86 pmol O_2_ min^-1^ µg^-1^) at any time point (Figure 3g). By contrast, *G. fascicularis* and *P. astreoides* exposed to hypoxia displayed significantly lower respiration rates (*G. fascicularis*: 37–52 pmol O_2_ min^-1^ µg^-1^; *P. astreoides*: 36–45 pmol O_2_ min^-1^ µg^-1^) compared to controls (*G. fascicularis*: 99–118 pmol O_2_ min^-1^ µg^-1^; *P. astreoides*: 92–102 pmol O_2_ min^-1^ µg^-1^) at all time points (Figure 3h–i).

### Effects of hypoxia on endosymbiont density and photophysiology

Exposure to hypoxia had significant, largely negative effects on the density and photophysiology of endosymbionts within *G. fascicularis* and *P. astreoides*. Photosynthesis rates (expressed as pmol O_2_ produced minute^-1^ µg AFDW^-1^) of *P. astreoides* larvae and juveniles were significantly depressed by hypoxia (Table S2). Specifically, photosynthesis rates for organisms exposed to hypoxia were significantly lower (14–29 pmol O_2_ min^-1^ µg^-1^) than those for controls (69–83 pmol O_2_ min^-1^ µg^-1^) at all time points (Figure 4a). Symbiont densities (expressed as symbiont cells µg AFDW^-1^) of *P. astreoides* larvae and juveniles were also significantly reduced by hypoxia (Table S2). Indeed, trends in this metric closely resembled those in photosynthesis rates, with organisms exposed to hypoxia displaying significantly lower endosymbiont densities (7–12 cells µg^-1^) compared to controls (17–24 cells µg^-1^) at all time points (Figure 4b). Hypoxia also hindered symbiosis establishment in *G. fascicularis* juveniles, with animals previously exposed to hypoxia displaying significantly lower symbiont densities (1 ± 1 cells µg^-1^) compared to controls (4 ± 1 cells µg^-1^) at 84 h (Table S2; Figure 4c). Similarly, the dark-adapted photochemical yield (F_v_/F_m_) of photosystem II in *P. astreoides* larval and juvenile symbiont was significantly inhibited by hypoxia (Table S2). As with photosynthesis rates and symbiont densities, photochemical yield was significantly lower for organisms exposed to hypoxia (0.025–0.266) compared to controls (0.339–0.452) at all time points (Figure 4d). Interestingly, trends in symbiont chlorophyll content (expressed as pg cell^-1^) did not match those observed for the other metrics, with symbionts residing within animals exposed to hypoxia displaying significantly higher chlorophyll content (20–76 pg cell^-1^) compared to controls (14–50 pg cell^-1^) at 12 and 36 h (Figure 4e).

**Figure 4.**
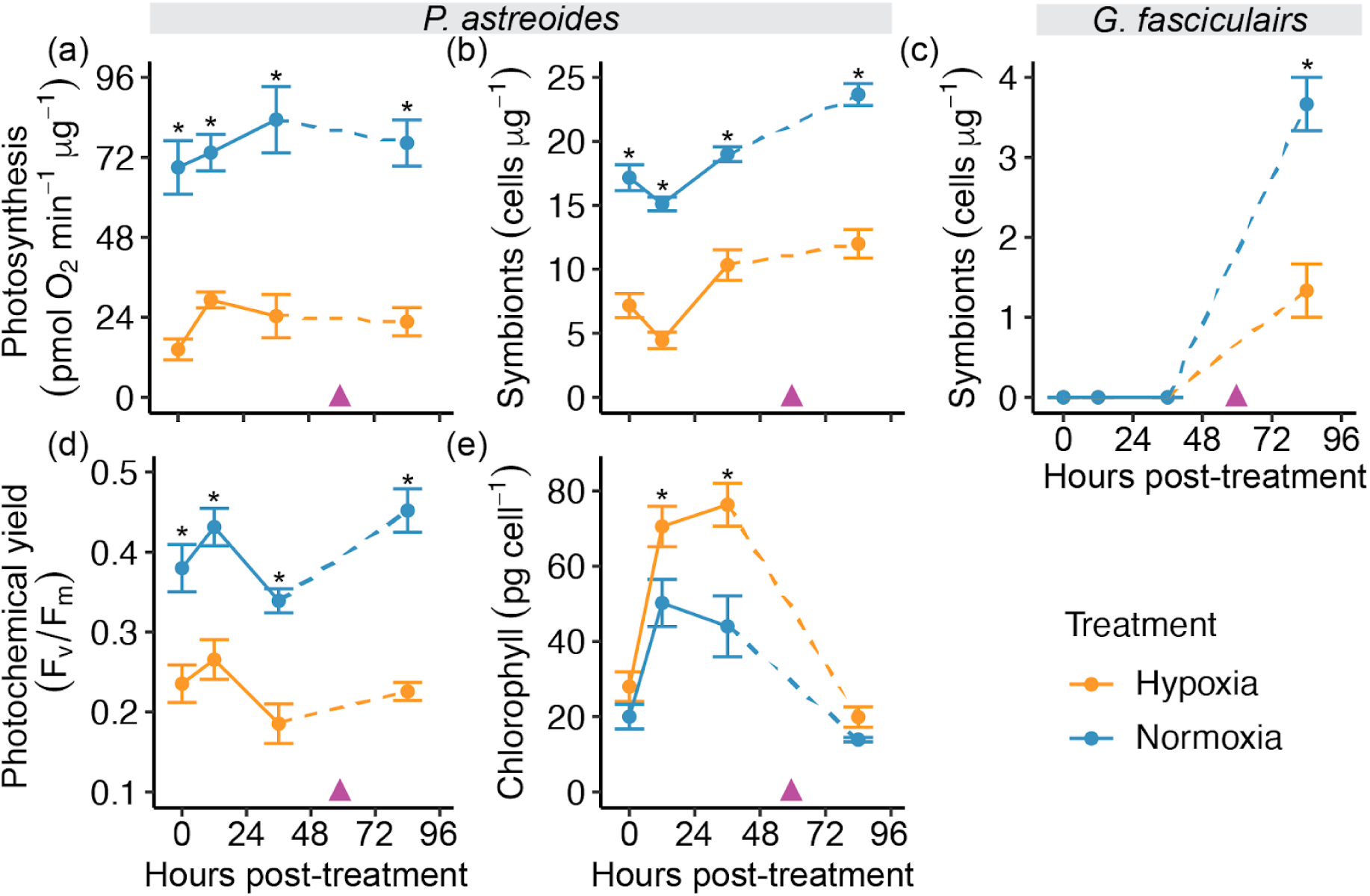
Effects of hypoxia on endosymbiont density and photophysiology. (a) Photosynthesis rates (pmol O_2_ produced min^-1^ µg AFDW^-1^) of *Porites astreoides* larval and juvenile (*n* = 60–90 treatment^-1^ time point^-1^ cohort^-1^ species^-1^; dashed line separates life stages) holobionts (i.e., animal and dinoflagellate endosymbionts together). (b–c) Symbiont density (cells µg AFDW^-1^) of (b) *P. astreoides* and (c) *Galaxea fascicularis* animals (*n* = 60–90 treatment^-1^ time point^-1^ cohort^-1^ species^-1^). (d) Photochemical yield (F_v_/F_m_) and (e) chlorophyll content (pg cell^-1^) of *P. astreoides* endosymbionts (*n* = 60–90 treatment^-1^ time point^-1^ cohort^-1^ species^-1^). All panels depict data over time following the hypoxia (yellow) and normoxia (blue) treatments. Purple arrowheads indicate the approximate timing of settlement, points with error bars depict means ± SEM, and asterisks indicate statistical significance (*p* < 0.05) of pairwise comparisons (hypoxia vs. normoxia).

### Effects of hypoxia on larval thermal tolerance

Upon being placed under acute heat stress at 36°C, larvae of all three species displayed progressive mortality over time (Figure 5a–l). Larval survival (expressed as a percentage) under acute heat stress was significantly influenced by several factors, including treatment and the interaction between species, h post-treatment, and h at 36°C (Table S2). For *N. vectensis* and *P. astreoides*, lethal doses 50 (LD50s; expressed in h) derived from survival curves were not significantly different for larvae previously exposed to hypoxia (*N. vectensis*: 78.5–84.5 h; *P. astreoides*: 44.2–50.8 h) compared to controls (*N. vectensis*: 81.2–87.8 h; *P. astreoides*: 45.2–52.1 h) at any time point (Figure 5a–c, g–i). By contrast, LD50s for *G. fascicularis* were significantly lower for larvae previously exposed to hypoxia at all time points (9.4–14.3 h) compared to controls (19.4–24.7 h) at the same time points (Figure 5d–f). In contrast to survival, LD50s (in h) for the decline in photochemical yield (F_v_/F_m_) of *P. astreoides* symbiont under acute heat stress were significantly depressed in animals previously exposed to hypoxia (15.2–18.3 h) compared to controls (21.6–23.4 h) at 12 and 36 h (Table S2).

**Figure 5.**
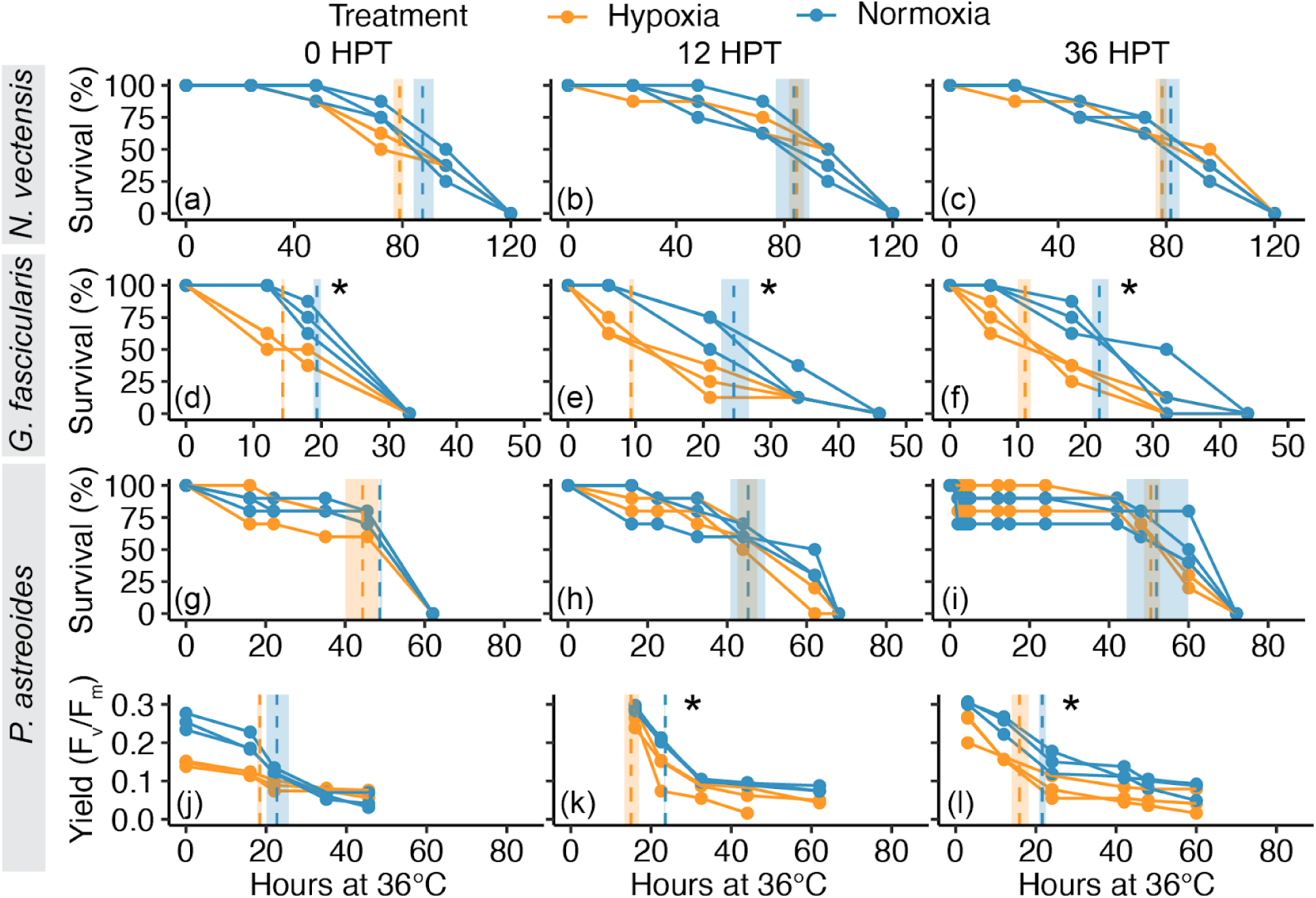
Effects of hypoxia on larval thermal tolerance. (a–i) Survival (%) of *Nematostella vectensis* (top), *Galaxea fascicularis* (middle), and *Porites astreoides* (bottom) larvae over time at 36°C for heat tolerance assays initiated at 0 (left), 12 (middle), and 36 (right) hours post-treatment (h; *n* = 30 larvae treatment^-1^ time point^-1^ cohort^-1^ species^-1^). (j–l) Photochemical yield (yield; F_v_/F_m_) of *P. astreoides* dinoflagellate endosymbionts over time at 36°C for heat tolerance assays initiated at 0 (left), 12 (middle), and 36 (right) h (*n* = 30 larvae treatment^-1^ time point^-1^ cohort^-1^ species^-1^). Data from larvae previously exposed to hypoxia are in yellow, while those from normoxia controls are in blue. Points represent raw data, while vertical dotted lines with shaded ribbons represent mean lethal doses 50 (LD50s) ± SEM. Asterisks indicate significance (*p* < 0.05) for pairwise comparisons of LD50s (hypoxia vs. normoxia).

## Discussion

Anthropogenic ocean deoxygenation presents a growing threat to global marine biodiversity (Altieri et al. 2017; Breitburg et al. 2018; Hughes et al. 2020; Rose et al. 2024). Early life stages of marine invertebrates are particularly sensitive to abiotic stressors (Przeslawski et al. 2015; Foo and Byrne 2017), making it critical to understand the impacts of stress exposure during these essential life history periods. Specifically, the effects of hypoxia on cnidarian early life stages are largely unknown, and it is unclear how evolutionary and life histories may influence these outcomes. Our results suggest numerous negative effects of hypoxia on the performance of cnidarian early life stages across a range of taxa. Thus, limiting further anthropogenic ocean deoxygenation and warming is vital for the persistence of cnidarian species along with the ecosystem functions and services they support.

### Reduced life cycle progression following hypoxia may threaten species persistence

*Nematostella vectensis*, *Galaxea fascicularis*, and *Porites astreoides* all exhibited declines in larval swimming behavior at the end of a 6-h hypoxia exposure. This behavioral change may limit larval dispersal, thereby decreasing connectivity and gene flow between habitats, and potentially contributing to increased competition for benthic space due to higher local recruitment (Baird et al. 2009). Reduced swimming may have benefitted larvae by temporarily decreasing their O_2_ demand and ATP consumption, permitting them to conserve energy until O_2_ availability increased. Indeed, significant decreases in respiration rates were observed in the coral larvae immediately following hypoxia exposure, although this was not true for *N. vectensis*, suggesting that reducing swimming behavior did not decrease respiratory activity in this species. Interestingly, larval survival and swimming appeared normal following the return to normoxic conditions for all three species, whereas respiration rates remained depressed in the corals through the juvenile stage. These results demonstrate that behavioral effects were transient while metabolic outcomes persisted. This persistent metabolic depression may promote larval fitness under hypoxia by decreasing O_2_ demand, which would be particularly beneficial if multiple hypoxic events occurred in rapid succession, as is sometimes observed in reef and estuary habitats (Summers 2001; Pezner et al. 2023). Metabolic depression may have resulted from damage to respiratory machinery (e.g., mitochondria) by reactive oxygen species (ROS), which are often produced following hypoxia-induced oxidative stress in corals and other organisms (Alderdice et al. 2021, 2022b; Zhang et al. 2023a). Larvae of other cnidarians, including the corals *Acropora cytherea* and *A. pulchra*, and the jellyfish *Aurelia aurita,* also display temporary reductions in normal swimming behavior following hypoxia exposure, suggesting shared behavioral responses to hypoxia that may be associated with conserved effects on metabolism (Jorissen and Nugues 2021). However, hypoxia does not affect larval swimming behavior in all coral species (Alderdice et al. 2022b; Mallon et al. 2023), indicating some species-specific differences in susceptibility that may influence the taxonomic composition of future reefs.

Declines in larval swimming behavior and metabolism during and after hypoxic events may have implications for ecological and evolutionary dynamics in reefs and estuaries due to downstream effects on settlement. Here, the two coral species experienced decreases in settlement rates following hypoxia exposure despite access to suitable substrates. Decreased settlement rates could have important consequences for the survival of populations exposed to hypoxic events, as generating new reproductive adults requires successful settlement. Potentially compounding these effects, hypoxia can also alter benthic community composition (Johnson et al. 2021a), thereby decreasing the abundance of biofilms and organisms that promote settlement (Cheung et al. 2014). The mechanisms resulting in decreased settlement are unknown, with one possibility being that hypoxia disrupted larval physiological processes required for attachment and metamorphosis (e.g., via the exhaustion of nutritional reserves). Additionally, hypoxia exposure can lead to the differential expression of genes involved in developmental regulation (e.g., homeobox [*HOX*] genes) in coral larvae and adults (Alderdice et al. 2021, 2022b), many of which are conserved targets of the transcription factor hypoxia-inducible factor (HIF). Intriguingly, *N. vectensis* larvae did not display differences in settlement following hypoxia exposure, demonstrating developmental resilience that may be an adaptation to the often extreme physicochemical conditions of the estuarine environments inhabited by this species (Summers 2001; Kiddon et al. 2003). Characterizing the molecular mechanisms underlying developmental outcomes following hypoxia exposure in cnidarians is an important future direction.

### Hypoxia influences size and growth, with important fitness implications

Changes in the size and growth of larvae and juveniles of all three species following hypoxia exposure suggest latent metabolic effects that may reduce fitness in future oceans. For example, larvae of *G. fascicularis* and *P. astreoides* displayed a persistent increase in body size following exposure to hypoxia that did not correspond with an increase in biomass. These results suggest increased cellular water content in larvae, and the mechanisms driving this response are unknown. One possible explanation could be an increase in cell/tissue necrosis (Helgoe et al. 2024), which can lead to the deactivation of cell membrane pumps and a subsequent influx of water, resulting in cell swelling (Khalid and Azimpouran 2023). Additionally, hypoxia exposure may have precipitated a switch to anaerobic respiration, as is observed in adult corals (Linsmayer et al. 2020). This could lead to the production of osmolytes (e.g., opines) that remain trapped in the cytoplasm of coral cells (Linsmayer et al. 2020), potentially causing osmotic swelling. Finally, the upregulation of aquaporins at the transcript level has been observed in larvae and adult corals following hypoxia exposure (Alderdice et al. 2022b), and might allow for compensatory O_2_ diffusion via aquaporin-mediated O_2_ influx accompanied by increased water uptake and swelling (Zwiazek et al. 2017). Aquaporins appear to play a conserved role in the response to hypoxia in a broad diversity of marine and terrestrial species including corals, exhibiting changes in expression, localization, and gating (Gobler et al. 2014; Alderdice et al. 2022b). However, *N. vectensis* larvae did not display the same increases in size observed in the coral species following hypoxia exposure, demonstrating taxonomically divergent responses and an interesting route for future research surrounding mechanisms of hypoxia resilience.

Surprisingly, the increases in size observed in larvae were reversed at the juvenile stage for all three species, with animals previously exposed to hypoxia exhibiting significantly smaller juvenile sizes than normoxia controls. These results indicate persistent costs incurred following hypoxia that reduced growth, which could have significant consequences for fitness. Specifically, smaller juveniles sizes are correlated with lower survival (Goodbody-Gringley et al. 2018), poorer competitive performance for space on the benthos (Tanner 1997), and lower fecundity upon reaching sexual maturity (Marshall and Keough 2007) in corals and other marine invertebrates. Indeed, larvae of the coral *Orbicella faveolata* exposed to hypoxia display reductions in juvenile survival (Mallon et al. 2023). It was surprising that *N. vectensis* juveniles exposed to hypoxia during the larval stage displayed smaller sizes than controls, particularly given that most of the other metrics quantified here were unaffected in this species. These findings imply that even transient hypoxic events can have lasting inhibitory effects on organismal performance.

### Hypoxia disrupts obligate coral-dinoflagellate mutualisms

Hypoxia destabilized the obligate coral-dinoflagellate mutualism during early life stages of both *G. fascicularis* and *P. astreoides*, indicating threatened holobiont success in future oceans. In *P. astreoides,* which inherits symbionts vertically and thus contained algae during the exposure period, hypoxia led to depressed photosynthesis rates, reduced photochemical yield, and symbiont loss (i.e., bleaching) that persisted through the juvenile stage. These results indicate lasting damage to symbiont photosynthetic machinery and the stability of this symbiosis, demonstrating that hypoxia can induce bleaching in larvae that persists through settlement. Importantly, the declines in symbiont density and performance in *P. astreoides* likely prevented larvae and juveniles of this species from acquiring sufficient photosynthate to meet their metabolic needs (Kitchen et al. 2020), perhaps contributing to the reductions in settlement and growth. The cause of these responses remains unknown and are an important topic for further investigation. One possibility is that hypoxia triggered ROS production in symbiont chloroplasts (Lee et al. 2023), which can depress photosynthesis rates in coral holobionts under hypoxia (Pontes et al. 2023). Unexpectedly, reduced symbiont performance and density were correlated with an increase in chlorophyll content per symbiont cell in *P. astreoides*, possibly representing an attempt to improve photosynthetic output. However, this increase in chlorophyll did not last through the juvenile stage, suggesting that some of the negative effects of hypoxia diminished over time. Symbiont photobiology and density is similarly affected by hypoxia in adult corals, contributing to mass bleaching and mortality during and after hypoxic events (Hughes et al. 2020; Johnson et al. 2021a; Swaminathan et al. 2024). Importantly, these results suggest that vertically transmitting coral species may be more threatened by hypoxia than horizontal transmitters, thus influencing the taxonomic composition of future reefs.

Interestingly, the coral *G. fascicularis,* which does not acquire symbionts until the juvenile stage, displayed disrupted symbiosis establishment following hypoxia exposure at the larval stage. These results suggest carryover effects on juvenile physiology that diminished symbiont uptake, establishment, and/or proliferation (Jacobovitz et al. 2023). The cause of this response is unclear, particularly since little is known regarding the molecular mechanisms regulating cnidarian symbiosis establishment (Jacobovitz et al. 2023). As the coral-dinoflagellate mutualism is obligate for shallow reef-building species (Helgoe et al. 2024), it is vital to further characterize the mechanisms by which hypoxia influences symbiosis establishment.

### Latent effects of hypoxia exposure exacerbated heat stress responses in corals

Hypoxia exposure led to poorer heat stress responses in the larvae of both corals, suggesting threatened performance under combined ocean deoxygenation and warming. These responses were more severe in *G. fascicularis*, where hypoxia-exposed individuals displayed lower survival under acute heat stress at all time points. While *P. astreoides* exposed to hypoxia did not display lower survival under heat stress relative to controls, they did exhibit faster declines in photochemical yield (F_v_/F_m_). These results indicate interactive effects of hypoxia and heat stress that may threaten the persistence of these species under combined ocean deoxygenation and warming. While the mechanisms of these effects are unknown, changes in the expression of heat stress response genes (e.g., heat shock proteins) coordinated by HIF have been observed in coral larvae and adults under hypoxia (Alderdice et al. 2022b; a). Notably, *N. vectensis* larvae exposed to hypoxia did not display lower survival under heat stress, again indicating robust stress resilience in this species that is likely a product of adaptation and (intergenerational) acclimatization to high DO and temperature variability in the estuarine environments inhabited by this species (Summers 2001; Rivera et al. 2021; Glass et al. 2023a). All together, these results suggest that coral larvae and their symbionts were sensitized to heat stress following hypoxia exposure. This sensitivity may undermine reef persistence, particularly since coral mass spawning can drive hypoxia and occurs during warm summer months when marine heatwaves and hypoxic events can coincide (Simpson et al. 1993; Pezner et al. 2023; Richards et al. 2024). However, transient exposure to stressors may also prime organisms for future exposure to the same stressor, a process known as hormetic priming, and it remains to be determined if hormetic priming to hypoxia occurs in cnidarians as it does for temperature and pH stress in corals and sea anemones (Putnam et al. 2020; Martell 2023; Glass et al. 2023a). Furthermore, cross-protection (i.e., the preparation for one stressor following exposure to a different stressor) has been observed in a broad diversity of species (Rodgers and Gomez Isaza 2021), and may also improve cnidarian resilience. The molecular mechanisms underpinning the effects of combined hypoxia and heat stress remain an important outstanding topic to further clarify how coral and sea anemone early life stages may fare in future seas.

### Conclusions

Here, we found that exposure to a simulated hypoxic event had myriad negative effects on larvae and juveniles of the estuarine sea anemone *N. vectensis* and reef-building corals *G. fascicularis* and *P. astreoides*. Comparing these three species revealed that a combination of evolved adaptations, (parental) environmental history, and life history traits all influence organismal hypoxia responses, suggesting differential success in future oceans. Specifically, *N. vectensis* displayed decreased larval swimming behavior immediately following hypoxia exposure, yet showed no changes in larval metabolic performance, heat tolerance, or settlement success, findings all in line with this being a stress-tolerant species (Darling et al. 2005; Reitzel et al. 2013; Friedman et al. 2018; Rivera et al. 2021; Glass et al. 2023b; a). This hypoxia resilience is likely an evolved adaptation to the transient hypoxia that occurs in this species’ native estuarine habitats (Summers 2001; Kiddon et al. 2003; Vaquer-Sunyer and Duarte 2011; Galic et al. 2019), though the exact mechanisms that support this are unclear and are an important topic of further investigation. Further, the adult population from which gametes were sourced may have been previously exposed to hypoxia in the wild (Summers 2001; Kiddon et al. 2003), suggesting that intergenerational plasticity via parental effects may also play a role in promoting hypoxia resilience as it does for heat (Rivera et al. 2021) and pH (Glass et al. 2023b) stress in this species, though this is also currently speculative and a key topic for future investigation. Importantly, *N. vectensis* juveniles previously exposed to hypoxia were significantly smaller than controls, suggesting that adaptations and intergenerational plasticity may be insufficient to fully protect this species from the negative effects of future ocean deoxygenation.

In contrast to *N. vectensis*, the corals displayed decreases in swimming behavior, metabolic performance, heat tolerance, and settlement success following hypoxia exposure, demonstrating a lack of adaptive mechanisms for dealing with this increasingly prevalent stressor. Interestingly, *G. fascicularis* larvae previously exposed to hypoxia were sensitized to heat stress, and while the symbiotic larvae of *P. astreoides* exposed to hypoxia did not display differences in survival under heat compared to controls, symbiont proliferation and function were reduced. These findings indicate that symbiotic coral larvae may be more susceptible to hypoxia compared to horizontally transmitting species — a hypothesis further supported by the significantly lower settlement rates of *P. astreoides* compared to *G. fascicularis* following hypoxia exposure. However, it was also interesting that growth at the juvenile stage was more strongly suppressed by previous hypoxia exposure in *N. vectensis* and *G. fascicularis*, whereas only *P. astreoides* juveniles displayed significant increases in sizes compared to larvae. This finding may reflect a benefit of larval symbiosis and/or increased hypoxia sensitivity of larvae from broadcast spawners compared to brooding species. Further research should investigate these hypotheses to clarify how life history traits determine the sensitivity of cnidarian early life stages to hypoxia. Overall, these findings demonstrate that even a single night of hypoxia can jeopardize cnidarian performance, thus emphasizing the importance of limiting further ocean deoxygenation for the stability of coastal marine biodiversity and ecosystem function.

## Acknowledgements

This work was supported by National Institutes of Health (NIH) Predoctoral T32 HD083185 to BHG, the American Fisheries Society Steven Berkeley Marine Conservation Fellowship to BHG, and National Science Foundation (NSF) awards 1923743 and 2237658 to KLB. Adult *Porites astreoides* colonies and larvae were collected under Protected Species Licence 2023070505, and samples were exported from Bermuda to the United States under Special Permit SP230802 and CITES export permit 23BM0007. The authors thank members of the Drs. Hollie Putnam, Samantha de Putron, Yvonne Sawall, and Phillip Cleves labs for assistance with the sourcing of coral larvae. HP, SP, and YS also acknowledge NSF award 2129274 and the Heising-Simons Foundation International award for project “Enhancing Coral Resilience.”

## Author contributions

Conceptualization: BHG, KLB; Data curation: KLB; Formal analysis: KLB; Funding acquisition: BHG, KLB; Investigation: BHG, KLB; Methodology: BHG, KLB; Project administration: BHG, KLB; Resources: BHG, KLB; Software: BHG, KLB; Supervision: KLB; Validation: BHG, KLB; Visualization: BHG, KLB; Writing – original draft: BHG, KLB; Writing – review & editing: BHG, KLB.

## Data availability statement

The data that support the findings of this study are openly available in Dryad at https://datadryad.org/stash/share/CvNhVXNKIprwRhjixnd-50FXCUc96FIvwON5jIIURSY.

## References

Alderdice, R., G. Perna, A. Cárdenas, and others. 2022a. Deoxygenation lowers the thermal threshold of coral bleaching. Sci. Rep. 12: 18273.

Alderdice, R., M. Pernice, A. Cárdenas, and others. 2022b. Hypoxia as a physiological cue and pathological stress for coral larvae. Mol. Ecol. 31: 571–587.

Alderdice, R., D. J. Suggett, A. Cárdenas, D. J. Hughes, M. Kühl, M. Pernice, and C. R. Voolstra. 2021. Divergent expression of hypoxia response systems under deoxygenation in reef-forming corals aligns with bleaching susceptibility. Glob. Chang. Biol. 27: 312–326.

Altieri, A. H., S. B. Harrison, J. Seemann, R. Collin, R. J. Diaz, and N. Knowlton. 2017. Tropical dead zones and mass mortalities on coral reefs. Proc. Natl. Acad. Sci. U. S. A. 114: 3660–3665.

Altieri, A. H., M. D. Johnson, S. D. Swaminathan, H. R. Nelson, and K. B. Gedan. 2021. Resilience of Tropical Ecosystems to Ocean Deoxygenation. Trends Ecol. Evol. 36: 227–238.

Alva García, J. V., S. G. Klein, T. Alamoudi, S. Arossa, A. J. Parry, A. Steckbauer, and C. M. Duarte. 2022. Thresholds of hypoxia of two Red Sea coral species (Porites sp. and Galaxea fascicularis). Frontiers in Marine Science 9. doi:10.3389/fmars.2022.945293

Baird, A. H., J. R. Guest, and B. L. Willis. 2009. Systematic and biogeographical patterns in the reproductive biology of scleractinian corals. Annu. Rev. Ecol. Evol. Syst. 40: 24.

Banse, K., and G. C. Anderson. 1967. Computations of chlorophyll concentrations from spectrophotometric readings. Limnol. Oceanogr. 12: 696–697.

Baumann, H., R. B. Wallace, T. Tagliaferri, and C. J. Gobler. 2015. Large Natural pH, CO2 and O2 Fluctuations in a Temperate Tidal Salt Marsh on Diel, Seasonal, and Interannual Time Scales. Estuaries Coasts 38: 220–231.

Breitburg, D., L. A. Levin, A. Oschlies, and others. 2018. Declining oxygen in the global ocean and coastal waters. Science 359. doi:10.1126/science.aam7240

Brown, K. T., E. A. Lenz, B. H. Glass, and others. 2023. Divergent bleaching and recovery trajectories in reef-building corals following a decade of successive marine heatwaves. Proc. Natl. Acad. Sci. U. S. A. 120: e2312104120.

Cheung, S. G., C. Y. S. Chan, B. H. K. Po, and others. 2014. Effects of hypoxia on biofilms and subsequently larval settlement of benthic invertebrates. Mar. Pollut. Bull. 85: 418–424.

Craggs, J., J. R. Guest, M. Davis, J. Simmons, E. Dashti, and M. Sweet. 2017. Inducing broadcast coral spawning ex situ: Closed system mesocosm design and husbandry protocol. Ecol. Evol. 7: 11066–11078.

Darling, J. A., A. R. Reitzel, P. M. Burton, M. E. Mazza, J. F. Ryan, J. C. Sullivan, and J. R. Finnerty. 2005. Rising starlet: the starlet sea anemone, Nematostella vectensis. Bioessays 27: 211–221.

DeCarlo, T. M., H. B. Harrison, L. Gajdzik, and others. 2019. Acclimatization of massive reef-building corals to consecutive heatwaves. Proc. Biol. Sci. 286: 20190235.

Dellaert, Z., and H. M. Putnam. 2023. Reconciling the variability in the biological response of marine invertebrates to climate change. J. Exp. Biol. 226. doi:10.1242/jeb.245834

Diaz, R. J., R. J. Neubauer, L. C. Schaffner, L. Pihl, and S. P. Baden. 1992. Continuous monitoring of dissolved oxygen in an estuary experiencing periodic hypoxia and the effect of hypoxia on macrobenthos and fish, p. 1055–1068. In R.A. Vollenweider, R. Marchetti, and R. Viviani [eds.], Marine Coastal Eutrophication. Elsevier.

Dilernia, N. J., S. Woodcock, E. F. Camp, D. J. Hughes, M. Kühl, and D. J. Suggett. 2024. Intra-colony spatial variance of oxyregulation and hypoxic thresholds for key Acropora coral species. Ecol. Evol. 14: e11100.

Foo, S. A., and M. Byrne. 2017. Marine gametes in a changing ocean: Impacts of climate change stressors on fecundity and the egg. Mar. Environ. Res. 128: 12–24.

Friedman, L. E., T. D. Gilmore, and J. R. Finnerty. 2018. Intraspecific variation in oxidative stress tolerance in a model cnidarian: Differences in peroxide sensitivity between and within populations of Nematostella vectensis. PLoS One 13: e0188265.

Galic, N., T. Hawkins, and V. E. Forbes. 2019. Adverse impacts of hypoxia on aquatic invertebrates: A meta-analysis. Sci. Total Environ. 652: 736–743.

Glass, B., and K. Barott. 2024. Data and code for: Hypoxia threatens coral and sea anemone early life stages.doi:10.5061/dryad.bnzs7h4hr

Glass, B. H., K. G. Jones, A. C. Ye, A. G. Dworetzky, and K. L. Barott. 2023a. Acute heat priming promotes short-term climate resilience of early life stages in a model sea anemone. PeerJ 11: e16574.

Glass, B. H., A. H. Schmitt, K. T. Brown, K. F. Speer, and K. L. Barott. 2023b. Parental exposure to ocean acidification impacts gamete production and physiology but not offspring performance in Nematostella vectensis. Biol. Open. doi:10.1242/bio.059746

Gobler, C. J., E. L. DePasquale, A. W. Griffith, and H. Baumann. 2014. Hypoxia and acidification have additive and synergistic negative effects on the growth, survival, and metamorphosis of early life stage bivalves. PLoS One 9: e83648.

Goodbody-Gringley, G., K. H. Wong, D. M. Becker, K. Glennon, and S. J. de Putron. 2018. Reproductive ecology and early life history traits of the brooding coral, Porites astreoides, from shallow to mesophotic zones. Coral Reefs 37: 483–494.

Hand, C., and K. R. Uhlinger. 1992. The Culture, Sexual and Asexual Reproduction, and Growth of the Sea Anemone Nematostella vectensis. Biol. Bull. 182: 169–176.

Harrison, P. L. 2011. Sexual Reproduction of Scleractinian Corals, p. 59–85. In Z. Dubinsky and N. Stambler [eds.], Coral Reefs: An Ecosystem in Transition. Springer Netherlands.

Helgoe, J., S. K. Davy, V. M. Weis, and M. Rodriguez-Lanetty. 2024. Triggers, cascades, and endpoints: connecting the dots of coral bleaching mechanisms. Biol. Rev. Camb. Philos. Soc. doi:10.1111/brv.13042

Hughes, D. J., R. Alderdice, C. Cooney, M. Kühl, M. Pernice, C. R. Voolstra, and D. J. Suggett. 2020. Coral reef survival under accelerating ocean deoxygenation. Nat. Clim. Chang. 10: 296–307.

Innis, T., L. Allen-Waller, K. T. Brown, and others. 2021. Marine heatwaves depress metabolic activity and impair cellular acid-base homeostasis in reef-building corals regardless of bleaching susceptibility. Glob. Chang. Biol. 27: 2728–2743.

Jacobovitz, M. R., E. A. Hambleton, and A. Guse. 2023. Unlocking the Complex Cell Biology of Coral-Dinoflagellate Symbiosis: A Model Systems Approach. Annu. Rev. Genet. doi:10.1146/annurev-genet-072320-125436

Johnson, M. D., J. J. Scott, M. Leray, N. Lucey, L. M. R. Bravo, W. L. Wied, and A. H. Altieri. 2021a. Rapid ecosystem-scale consequences of acute deoxygenation on a Caribbean coral reef. Nat. Commun. 12: 4522.

Johnson, M. D., S. D. Swaminathan, E. N. Nixon, V. J. Paul, and A. H. Altieri. 2021b. Differential susceptibility of reef-building corals to deoxygenation reveals remarkable hypoxia tolerance. Sci. Rep. 11: 23168.

Jorissen, H., and M. M. Nugues. 2021. Coral larvae avoid substratum exploration and settlement in low-oxygen environments. Coral Reefs 40: 31–39.

Khalid, N., and M. Azimpouran. 2023. Necrosis, StatPearls Publishing.

Kiddon, J. A., J. F. Paul, H. W. Buffum, C. S. Strobel, S. S. Hale, D. Cobb, and B. S. Brown. 2003. Ecological condition of US Mid-Atlantic estuaries, 1997-1998. Mar. Pollut. Bull. 46: 1224–1244.

Kitchen, R. M., M. Piscetta, M. R. de Souza, E. A. Lenz, D. W. H. Schar, R. D. Gates, and C. B. Wall. 2020. Symbiont transmission and reproductive mode influence responses of three Hawaiian coral larvae to elevated temperature and nutrients. Coral Reefs 39: 419–431.

Krediet, C. J., J. C. DeNofrio, C. Caruso, M. S. Burriesci, K. Cella, and J. R. Pringle. 2015. Rapid, precise, and accurate counts of Symbiodinium cells using the Guava flow cytometer, and a comparison to other methods. PLoS One 10: e0135725.

Kvitt, H., A. Malik, S. Ben-Tabou de-Leon, and others. 2022. Transcriptional responses indicate acclimation to prolonged deoxygenation in the coral Stylophora pistillata. Front. Mar. Sci. 9. doi:10.3389/fmars.2022.999558

Lee, Y., E. Byeon, D.-H. Kim, and others. 2023. Hypoxia in aquatic invertebrates: Occurrence and phenotypic and molecular responses. Aquat. Toxicol. 263: 106685.

Linsmayer, L. B., D. D. Deheyn, L. Tomanek, and M. Tresguerres. 2020. Dynamic regulation of coral energy metabolism throughout the diel cycle. Sci. Rep. 10: 19881.

Linsmayer, L. B., S. K. Noel, M. Leray, D. Wangpraseurt, C. Hassibi, D. I. Kline, and M. Tresguerres. 2024. Effects of bleaching on oxygen dynamics and energy metabolism of two Caribbean coral species. Sci. Total Environ. 919: 170753.

Malcolm, J. M., and W. I. Brown. 1977. Zooxanthellae-produced O_2_ promotes sea anemone expansion and eliminates oxygen debt under environmental hypoxia. J. Exp. Zool. 201: 149–155.

Mallon, J. E., A. M. Demko, J. M. Sneed, L. Newman, C. Dugan, A. H. Altieri, V. J. Paul, and M. D. Johnson. 2023. The influence of deoxygenation on Caribbean coral larval settlement and early survival. Frontiers in Marine Science 10. doi:10.3389/fmars.2023.1254965

Marshall, D. J., and M. J. Keough. 2007. The Evolutionary Ecology of Offspring Size in Marine Invertebrates, p. 1–60. In Advances in Marine Biology. Academic Press.

Martell, H. A. 2023. Thermal priming and bleaching hormesis in the staghorn coral, Acropora cervicornis (Lamarck 1816). J. Exp. Mar. Bio. Ecol. 560: 151820.

Murphy, J. W. A., and R. H. Richmond. 2016. Changes to coral health and metabolic activity under oxygen deprivation. PeerJ 4: e1956.

Nelson, H. R., and A. H. Altieri. 2019. Oxygen: the universal currency on coral reefs. Coral Reefs 38: 177–198.

Pezner, A. K., T. A. Courtney, H. C. Barkley, and others. 2023. Increasing hypoxia on global coral reefs under ocean warming. Nat. Clim. Chang. 1–7.

Pontes, E., C. Langdon, and F. A. Al-Horani. 2023. Caribbean scleractinian corals exhibit highly variable tolerances to acute hypoxia. Frontiers in Marine Science 10. doi:10.3389/fmars.2023.1120262

Przeslawski, R., M. Byrne, and C. Mellin. 2015. A review and meta-analysis of the effects of multiple abiotic stressors on marine embryos and larvae. Glob. Chang. Biol. 21: 2122–2140.

Putnam, H. M., R. Ritson-Williams, J. A. Cruz, J. M. Davidson, and R. D. Gates. 2020. Environmentally-induced parental or developmental conditioning influences coral offspring ecological performance. Sci. Rep. 10: 13664.

de Putron, S. J., J. M. Lawson, K. Q. L. White, M. T. Costa, M. V. B. Geronimus, and A. MacCarthy. 2017. Variation in larval properties of the Atlantic brooding coral Porites astreoides between different reef sites in Bermuda. Coral Reefs 36: 383–393.

Rands, M. L., A. E. Douglas, B. C. Loughman, and R. G. Ratcliffe. 1992. Avoidance of Hypoxia in a Cnidarian Symbiosis by Algal Photosynthetic Oxygen. Biol. Bull. 182: 159–162.

Ranke, J. 2022. chemCal: Calibration Functions for Analytical Chemistry,.

Reitzel, A. M., T. Chu, S. Edquist, C. Genovese, C. Church, A. M. Tarrant, and J. R. Finnerty. 2013. Physiological and developmental responses to temperature by the sea anemone Nematostella vectensis. Mar. Ecol. Prog. Ser. 484: 115–130.

Ricardo, G. F., R. J. Jones, P. L. Clode, and A. P. Negri. 2016. Mucous Secretion and Cilia Beating Defend Developing Coral Larvae from Suspended Sediments. PLoS One 11: e0162743.

Richards, Z. T., L. Haines, C. Ross, and others. 2024. Deoxygenation following coral spawning and low-level thermal stress trigger mass coral mortality at Coral Bay, Ningaloo Reef. Coral Reefs. doi:10.1007/s00338-024-02476-x

Ritz, C., F. Baty, J. C. Streibig, and D. Gerhard. 2015. Dose-Response Analysis Using R. PLoS One 10: e0146021.

Rivera, H. E., C.-Y. Chen, M. C. Gibson, and A. M. Tarrant. 2021. Plasticity in parental effects confers rapid larval thermal tolerance in the estuarine anemone Nematostella vectensis. J. Exp. Biol. 224. doi:10.1242/jeb.236745

Rodgers, E. M., and D. F. Gomez Isaza. 2021. Harnessing the potential of cross-protection stressor interactions for conservation: a review. Conserv Physiol 9: coab037.

Rose, K. C., E. M. Ferrer, S. R. Carpenter, and others. 2024. Aquatic deoxygenation as a planetary boundary and key regulator of Earth system stability. Nat Ecol Evol. doi:10.1038/s41559-024-02448-y

Rosenberg, R. 1980. Effect of oxygen deficiency on benthic macrofauna in fjords, p. 499–514. In Fjord Oceanography. Springer US.

RStudio Team. 2020. RStudio: Integrated Development for R,.

Schindelin, J., I. Arganda-Carreras, E. Frise, and others. 2012. Fiji: an open-source platform for biological-image analysis. Nat. Methods 9: 676–682.

Shick, J. M. 1990. Diffusion Limitation and Hyperoxic Enhancement of Oxygen Consumption in Zooxanthellate Sea Anemones, Zoanthids, and Corals. Biol. Bull. 179: 148–158.

Simpson, C. J., J. L. Cary, and R. J. Masini. 1993. Destruction of corals and other reef animals by coral spawn slicks on Ningaloo Reef, Western Australia. Coral Reefs 12: 185–191.

Summers, J. K. 2001. Ecological condition of the estuaries of the Atlantic and Gulf Coasts of the United States. Environ. Toxicol. Chem. 20: 99–106.

Swaminathan, S. D., J. L. Meyer, M. D. Johnson, V. J. Paul, E. Bartels, and A. H. Altieri. 2024. Divergent responses of the coral holobiont to deoxygenation and prior environmental stress. Frontiers in Marine Science 10. doi:10.3389/fmars.2023.1301474

Tanner, J. E. 1997. Interspecific competition reduces fitness in scleractinian corals. J. Exp. Mar. Bio. Ecol.

Vaquer-Sunyer, R., and C. M. Duarte. 2011. Temperature effects on oxygen thresholds for hypoxia in marine benthic organisms. Glob. Chang. Biol. 17: 1788–1797.

Wangpraseurt, D., M. Weber, H. Røy, L. Polerecky, D. de Beer, Suharsono, and M. M. Nugues. 2012. In situ oxygen dynamics in coral-algal interactions. PLoS One 7: e31192.

Wei, F., M. Cui, W. Huang, Y. Wang, X. Liu, X. Zeng, H. Su, and K. Yu. 2023. Ex situ reproduction and recruitment of scleractinian coral Galaxea fascicularis. Mar. Biol. 170: 30.

Whitaker, J. R., and P. E. Granum. 1980. An absolute method for protein determination based on difference in absorbance at 235 and 280 nm. Anal. Biochem. 109: 156–159.

Wong, K. H., G. Goodbody-Gringley, S. J. de Putron, D. M. Becker, A. Chequer, and H. M. Putnam. 2021. Brooded coral offspring physiology depends on the combined effects of parental press and pulse thermal history. Glob. Chang. Biol. 27: 3179–3195.

Zhang, K., Z. Wu, Z. Liu, J. Tang, W. Cai, M. An, and Z. Zhou. 2023a. Acute hypoxia induces reduction of algal symbiont density and suppression of energy metabolism in the scleractinian coral Pocillopora damicornis. Mar. Pollut. Bull. 191: 114897.

Zhang, Y., S. E. Gantt, E. F. Keister, and others. 2023b. Performance of *Orbicella faveolata* larval cohorts does not align with previously observed thermal tolerance of adult source populations. Glob. Chang. Biol. doi:10.1111/gcb.16977

Zwiazek, J. J., H. Xu, X. Tan, A. Navarro-Ródenas, and A. Morte. 2017. Significance of oxygen transport through aquaporins. Sci. Rep. 7: 40411.

