## supplementary material for "Hypoxia threatens coral and sea anemone early life stages"

### Supplementary materials and methods

*Adult culture and spawning*

Adult *Nematostella vectensis* sea anemones were collected from a salt marsh in Brigantine, New Jersey in the fall of 2020. Following transport to the University of Pennsylvania, anemones were kept in 12 parts per thousand (ppt) artificial seawater (Spectrum Brands) at 18°C in a dark incubator (Boekel Scientific). Animals were fed twice per week with *Artemia* nauplii (Brine Shrimp Direct), with water changes occurring approximately every 2 weeks for ~3 years to remove waste and maintain seawater normoxia (> 6.5 mg dissolved oxygen (DO) $L^{-1}$). Spawning was induced using a standard method for *N. vectensis* (Hand and Uhlinger 1992; Fritzenwanker and Technau 2002; Stefanik et al. 2013), which entailed exposing anemones ($n$ = 200 adults across four containers) to light and elevated temperatures (24°C) for 14 h, followed by transfer to room temperature (~18–19°C), after which spawning occurred within 1–2 h. As culture containers housed both male and female anemones, gametes were left to fertilize following spawning and the resulting embryos were then transferred to a plastic dish with ~25 mL of new seawater. Embryos were placed at 18°C in the dark, where they were left for 3 days to develop into swimming planulae for use in the experiment. These procedures yielded a single cohort of larvae with mixed parentage.

A captive aquarium population of adult *Galaxea fascicularis* colonies located at Carnegie Science in Baltimore, MD, USA (*n* = 9 females, 10 males; symbiont community unknown) were spawned in an *ex situ* coral spawning system designed as previously described (Craggs et al. 2017, 2020; O'Neil et al. 2021). The colonies were maintained in aquaria under lights with the timing of sunset, sunrise, moon set, moon rise, and photoperiod mimicking those in Cairns, Australia (Craggs et al. 2017). The seawater temperature in the aquaria was maintained at a non-sequential 10-year average for this region derived from published data (Puntin et al. 2022). Spawning occurred over several days in August 2023. On the second day of spawning, bundles containing gametes were collected using a transfer pipette immediately upon release from parent colonies and placed in 50 mL conical tubes. Upon bundle dissociation, the eggs were washed with artificial seawater, and then fertilization was performed by combining eggs and sperm from multiple colonies into a single pool. These procedures yielded a cohort of larvae with mixed parentage, which were kept in an incubator (Percival) at 27°C under lights (25 µmol photons $m^{-2}$ $s^{-1}$) with a 12-h:12-h day:night cycle until experimentation.

Five days prior to the July 2023 new moon, adult *Porites astreoides* colones (*n* = 20; symbiont community unknown) with a surface area of ~170 $cm^2$ were collected via hammer and chisel at a depth of ~2–5 m from Bailey's Bay Flats (32°22'13" N, 64°44'27" W), a patch reef off the northern coast of Bermuda. Colonies were transported in coolers containing seawater to the Bermuda Marine Mesocosm Facility (BMMF) at the Bermuda Institute of Ocean Sciences, where they were then placed in individual kitchen jugs set atop egg crate in an outdoor tank (~400 L) with flow-through seawater. The temperature in the tank was controlled by a 300 W chiller (AccuTherm) set to recapitulate ambient temperatures (~27.5–29°C) at Bailey's Bay. To mimic natural irradiance conditions on the reef (Wong et al. 2021), the tank was covered with a screen of black mesh (500 µm), and water levels were maintained at ~50 cm above the colonies during the day. Larvae (endosymbiont community unknown) were collected according to previously developed procedures for this species and facility (Rivera and Goodbody-Gringley 2014; Reich et al. 2017; Goodbody-Gringley et al. 2018). Specifically, each night following the collection of the adult colonies, the water level in the holding tank was lowered to a point just below the lips of the containers, and tubes with lightly flowing seawater were placed alongside each colony in the containers. This setup ensured that any planulae released overnight would be carried by flowing seawater down the jug handles and into a mesh (150 µm) container set at the base of each handle. On four separate occasions (i.e., four larval cohorts), larvae collected overnight were pooled across the parental colonies and kept in one of the mesh collection containers in the tank for ~12 h prior to the experiment. The hypoxia experiment was repeated for each of the four cohorts. For all 3 species, the conditions under which larvae were cultured before and after the treatment period represent ambient conditions and match those experienced by the adult populations from which gametes were sourced. As each of these species requires abiotic conditions (e.g., temperature and salinity) not shared with the others, the larvae were not cultured under identical conditions, which may have influenced their hypoxia responses.

*Symbiosis establishment trials for* Galaxea fascicularis

To assay symbiosis establishment in aposymbiotic *G. fascicularis* juveniles, dinoflagellate endosymbionts (family Symbiodiniaceae; community composition unknown) were introduced into culture containers shortly after 84 h, at which time juveniles of this species are likely to be competent for symbiont uptake (Wei et al. 2023). To ensure taxonomic diversity and symbiotic amenability of supplied symbionts, *in hospite* endosymbionts were sourced from fragments of several colonies of the coral *Pocillopora acuta* and polyps of the sea anemone *Exaiptasia diaphana* kept in a laboratory aquarium. Specifically, *P. acuta* fragments were waterpikked with sterile seawater and the resulting tissue slurry was collected in a 50 mL conical. Next, two *E. diaphana* polyps were added to the tissue slurry and immediately homogenized using a rotostator at 25,000 rpm for 15 sec. The isolation of symbionts from host (i.e., coral and sea anemone) cells was confirmed via brightfield microscopy. The isolated symbionts were pelleted by centrifugation at 1,700 rpm for 5 min, then resuspended in 6 mL of seawater (estimated concentration: ~500,000 cells mL$^{-1}$), of which 1 mL was added to each culture container. After 6 h, water in culture containers was gently stirred to encourage the resuspension of settled symbionts. After an additional 18 h, a water change was performed to wash out free symbiont cells, and the symbiotic juveniles were sampled ($n$ = 15–30 juveniles treatment$^{-1}$) by culture container and stored at -80°C for later analysis of symbiont density.

*Data and statistical analyses*

RStudio with R version 4.2.1 was used for all analyses (RStudio Team 2020). Preliminary analyses suggested a lack of significant differences between the four *P. astreoides* cohorts, so cohort was not included as a variable in the following analyses. First, survival data from larval heat tolerance assays were used to create dose-response curves represented by two-parameter log-logistic functions using the package *drc* (Ritz et al. 2015), and an LD50 was determined for each curve ($n$ = 3 curves treatment$^{-1}$ time point$^{-1}$ cohort$^{-1}$ species$^{-1}$) using the package *chemCal* (Ranke 2022). For photochemical yield data from *P. astreoides* larval heat tolerance assays, dose-response curves represented by log-logistic functions were created also using *drc*, and LD50s were again determined for each curve ($n$= 3 curves treatment$^{-1}$ time point$^{-1}$ cohort$^{-1}$) using *chemCal*. Next, for data pertaining to all metrics (swimming behavior, settlement rates, size, ash-free dry weight, respiration, photosynthesis, photochemical yield, symbiont density, chlorophyll per symbiont, and LD50s), linear models were constructed relating each metric to the interaction between species, treatment, and/or time (h post-treatment) as appropriate. Group within each experimental treatment was included as a random effect variable in the linear models where appropriate and permissible by the data structure. Each of these linear models was confirmed to meet relevant assumptions (linearity, statistical independence of errors, homoscedasticity of errors, and normality of the error distribution) via visual inspection of diagnostic plots (residuals vs. fitted values, residuals vs. leverage, scale-location, and normal Q-Q, respectively) generated using the "plot(lm)" function. Next, each model was subjected to an analysis of variance (ANOVA) with type II (no interactive effects in model) or III (interactive effects present in model) sums of squares to determine the statistical significance of the model terms. Finally, Tukey's Honest Significant Difference post-hoc tests were used to determine the significance of pairwise comparisons where appropriate. Additional packages used for analysis include: *ggplot2* (Wickham 2016), *emmeans* (Lenth et al. 2023), *ggpubr* (Kassambara 2023), and *lemon* (Edwards 2023). All values are expressed as averages rounded to appropriate significant figures ± standard error of the mean (SEM), and all original data and code are publicly available online (Glass and Barott 2024).

### Supplementary tables and figures

**Table S1. Treatment seawater conditions (mean ± SEM).**

| Species | Treatment | Time in treatment (h) | Temperature (°C) | Dissolved oxygen (mg $L^{-1}$) | Dissolved oxygen (%) |
|---|---|---|---|---|---|
| *Nematostella vectensis* | Normoxia | 0 | 20.3 ± 0.1 | 8.69 ± 0 | 96.1 ± 0 |
| | Normoxia | 6 | 18 ± 0.1 | 8.67 ± 0 | 91.5 ± 0 |
| | Hypoxia | 0 | 20.2 ± 0.1 | 1.6 ± 0 | 17.7 ± 0 |
| | Hypoxia | 6 | 18.1 ± 0.1 | 1.58 ± 0 | 16.7 ± 0 |
| *Galaxea fascicularis* | Normoxia | 0 | 26 ± 0.1 | 7.59 ± 0 | 93.6 ± 0 |
| | Normoxia | 6 | 27.1 ± 0.1 | 7.58 ± 0 | 95.3 ± 0 |
| | Hypoxia | 0 | 25.8 ± 0.1 | 1.6 ± 0 | 19.7 ± 0 |
| | Hypoxia | 6 | 27.1 ± 0.1 | 1.7 ± 0 | 21.4 ± 0 |
| *Porites astreoides* | Normoxia | 0 | 27.4 ± 0.2 | 6.8 ± 0.1 | 86 ± 1.3 |
| | Normoxia | 6 | 28.1 ± 0.2 | 6.8 ± 0.1 | 87.1 ± 1.3 |
| | Hypoxia | 0 | 27.5 ± 0.2 | 1.6 ± 0 | 20.3 ± 0 |
| | Hypoxia | 6 | 28.1 ± 0.2 | 1.8 ± 0.2 | 23.1 ± 2.5 |

**Table S2. Statistical testing information.**

| **Metric** | **Degrees of freedom** | ***F* value (ANOVA)** | ***P* value (ANOVA)** | **Pairwise *p* values (Tukey's HSD)** |
|---|---|---|---|---|
| Swimming (%) | 2 | Species*treatment: 4.9246 | Species*treatment: 0.02 | Treatment: < 0.05 for all species |
| Settlement (%) | 2 | Species*treatment: 9.472 | Species*treatment: 0.001 | Treatment: 0.749 (*N. vectensis*; < 0.001 (others) |
| Size (larval length) | 6 | Species*treatment: 2.427 | Species*treatment: 0.024 | Treatment: > 0.05 (*N. vectensis*); < 0.05 (others) |
| Size (juvenile polyp diameter) | 6 | Species*treatment: 2.427 | Species*treatment: 0.024 | Treatment: < 0.05 (all) |
| Ash-free dry weight | 2 | Species*treatment: 5.2247 | Species*treatment: 0.007 | Treatment: > 0.05 (*N. vectensis*, *G. fascicularis* juveniles); < |

| | | | | |
|---|---|---|---|---|
| | | | | 0.05 (*G. fascicularis* larvae and *P. astreoides*) |
| Respiration | 2 | Species*treatment: 23.2474 | Species*treatment: < 0.001 | Treatment: > 0.05 (*N. vectensis*); < 0.05 (*G. fascicularis* and *P. astreoides*) |
| Photosynthesis | 1 | Treatment: 50.5192 | Treatment: < 0.001 | Treatment: < 0.001 (all time points) |
| Endosymbiont density (*P. astreoides*) | 1 | Treatment: 52.7837 | Treatment: < 0.001 | Treatment: < 0.001 (all time points) |
| Endosymbiont density (*G. fascicularis*) | 1 | Treatment*Hours post-treatment: 24.5 | Treatment: < 0.001 | Treatment: > 0.05 (larvae); < 0.001(juveniles) |
| Photochemical yield ($F_v/F_m$) | 1 | Treatment: 16.1945 | Treatment: < 0.001 | Treatment: < 0.05 (all time |

| | | | | |
|---|---|---|---|---|
| | | | | points) |
| Chlorophyll | 1 | Treatment*Hours post-treatment: 36.3112 | Treatment*Hours post-treatment: < 0.001 | Treatment: 0.2622 (0 h); 0.001 (12); 0.002 (36); 0.3025 (84) |
| Heat tolerance (survival) | 1 | Treatment: 4.3941 | Treatment: 0.037 | Treatment: > 0.05 (*N. vectensis* and *P. astreoides*); < 0.05 (*G. fascicularis*) |
| Heat tolerance ($F_v/F_m$) | 2 | Treatment*hours post-treatment*hours at 36°C: 3.6097 | Treatment*hours post-treatment*hours at 36°C: 0.031 | Treatment: 0.078 (0 h); 0.004 (12); 0.035 (36) |

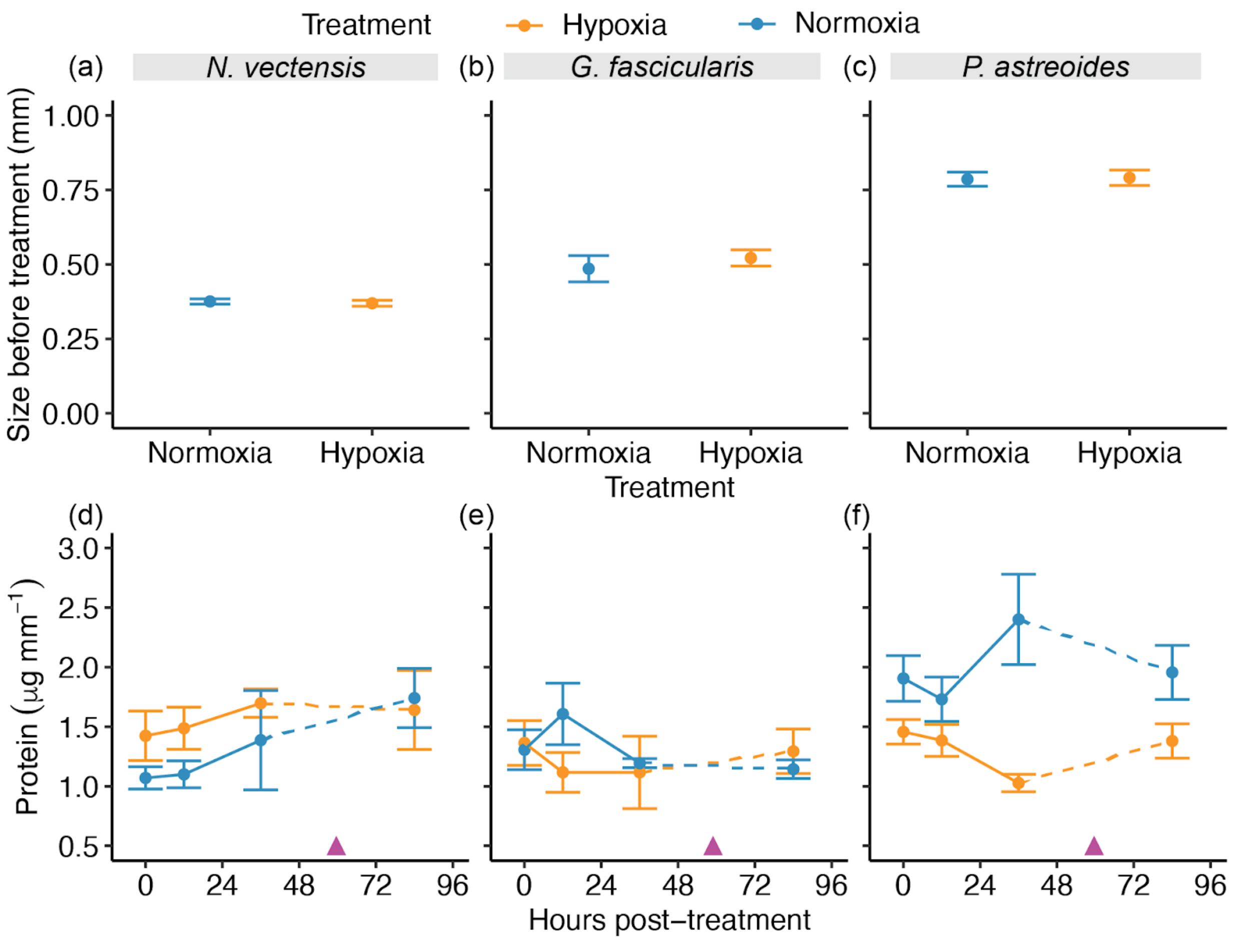


**Figure S1. Pre-treatment larval size and effects of hypoxia on protein.** (a–c) Size (larval length in mm) of *Nematostella vectensis* (left), *Galaxea fascicularis* (middle), and *Porites astreoides* (right) larvae in the hypoxia (yellow) and normoxia (blue) treatment groups ($n$ = 20–30 larvae treatment$^{-1}$ time point$^{-1}$ cohort$^{-1}$ species$^{-1}$) prior to the start of the experiment. (d–f) Protein (normalized to size; µg mm$^{-1}$) of *N. vectensis* (left), *G. fascicularis* (middle), and *P. astreoides* (right) larvae and juveniles ($n$ = 20–30 larvae treatment$^{-1}$ time point$^{-1}$ cohort$^{-1}$ species$^{-1}$) over time following the hypoxia (yellow) and normoxia (blue) treatments. In (d–f), purple arrowheads indicate the approximate timing of settlement, points with error bars depict means ± SEM, and asterisks indicate statistical significance ($p < 0.05$) of pairwise comparisons (hypoxia vs. normoxia).

### References


Craggs, J., J. Guest, M. Davis, and M. Sweet. 2020. Completing the life cycle of a broadcast spawning coral in a closed mesocosm. Invertebr. Reprod. Dev.

Craggs, J., J. R. Guest, M. Davis, J. Simmons, E. Dashti, and M. Sweet. 2017. Inducing broadcast coral spawning ex situ: Closed system mesocosm design and husbandry protocol. Ecol. Evol. **7**: 11066–11078.

Edwards, S. M. 2023. Lemon. Comprehensive R Archive Network (CRAN).

Fritzenwanker, J. H., and U. Technau. 2002. Induction of gametogenesis in the basal cnidarian Nematostella vectensis(Anthozoa). Dev. Genes Evol. **212**: 99–103.

Glass, B., and K. Barott. 2024. Data and code for: Hypoxia threatens coral and sea anemone early life stages.doi:10.5061/dryad.bnzs7h4hr

Goodbody-Gringley, G., K. H. Wong, D. M. Becker, K. Glennon, and S. J. de Putron. 2018. Reproductive ecology and early life history traits of the brooding coral, Porites astreoides, from shallow to mesophotic zones. Coral Reefs **37**: 483–494.

Hand, C., and K. R. Uhlinger. 1992. The Culture, Sexual and Asexual Reproduction, and Growth of the Sea Anemone Nematostella vectensis. Biol. Bull. **182**: 169–176.

Kassambara, A. 2023. ggpubr, Github.

Lenth, R. V., B. Bolker, P. Buerkner, and others. 2023. emmeans: Estimated marginal means, Github.

O'Neil, K. L., R. M. Serafin, and J. T. Patterson. 2021. Repeated ex situ Spawning in Two Highly Disease Susceptible Corals in the Family Meandrinidae. Frontiers in Marine.

Puntin, G., J. Craggs, R. Hayden, K. E. Engelhardt, S. McIlroy, M. Sweet, D. M. Baker, and M.

Ziegler. 2022. The reef-building coral Galaxea fascicularis: a new model system for coral symbiosis research. Coral Reefs. doi:10.1007/s00338-022-02334-8

Ranke, J. 2022. chemCal: Calibration Functions for Analytical Chemistry,.

Reich, H. G., D. L. Robertson, and G. Goodbody-Gringley. 2017. Do the shuffle: Changes in Symbiodinium consortia throughout juvenile coral development. PLoS One **12**: e0171768.

Ritz, C., F. Baty, J. C. Streibig, and D. Gerhard. 2015. Dose-Response Analysis Using R. PLoS One **10**: e0146021.

Rivera, H. E., and G. Goodbody-Gringley. 2014. Aggregation and cnidae development as early defensive strategies in Favia fragum and Porites astreoides. Coral Reefs **33**: 1079–1084.

RStudio Team. 2020. RStudio: Integrated Development for R,.

Stefanik, D. J., L. E. Friedman, and J. R. Finnerty. 2013. Collecting, rearing, spawning and inducing regeneration of the starlet sea anemone, Nematostella vectensis. Nat. Protoc. **8**: 916–923.

Wei, F., M. Cui, W. Huang, Y. Wang, X. Liu, X. Zeng, H. Su, and K. Yu. 2023. Ex situ reproduction and recruitment of scleractinian coral Galaxea fascicularis. Mar. Biol. **170**: 30.

Wickham, H. 2016. ggplot2: Elegant Graphics for Data Analysis, Springer-Verlag New York.

Wong, K. H., G. Goodbody-Gringley, S. J. de Putron, D. M. Becker, A. Chequer, and H. M. Putnam. 2021. Brooded coral offspring physiology depends on the combined effects of parental press and pulse thermal history. Glob. Chang. Biol. **27**: 3179–3195.
